# Amoeboid-Mesenchymal Transition and the Proteolytic Control of Cancer Invasion Plasticity

**DOI:** 10.1101/2025.10.24.684403

**Authors:** Adam W. Olson, Jonathan Li, Xiao-Yan Li, Lana King, Kalins Banerjee, Atticus J. McCoy, Mahnoor N. Gondal, Arul M. Chinnaiyan, Dorraya El-Ashry, Evan T. Keller, Andrew J. Putnam, Stephen J. Weiss

## Abstract

Invasion plasticity allows malignant cells to toggle between collective, mesenchymal and amoeboid phenotypes while traversing extracellular matrix (ECM) barriers. Current dogma holds that collective and mesenchymal invasion programs trigger the mobilization of proteinases that digest structural barriers dominated by type I collagen, while amoeboid activity allows cancer cells to marshal mechanical forces to traverse tissues independently of ECM proteolysis. Here, we use cancer spheroid-3-dimensional matrix models, single-cell RNA sequencing, and human tissue explants to identify the mechanisms controlling mesenchymal versus amoeboid invasion. Unexpectedly, collective/mesenchymal- and amoeboid-type invasion programs – though distinct – are each characterized by active tunneling through ECM barriers, with expression of matrix-degradative metalloproteinases. CRISPR/Cas9-mediated targeting of a single membrane-anchored collagenase, MMP14/MT1-MMP, ablates tissue-invasive activity while co-regulating cancer cell transcriptional programs. Though changes in matrix architecture, nuclear rigidity, and metabolic stress as well as the presence of cancer-associated fibroblasts are proposed to support amoeboid activity, none of these changes restore invasive activity of MMP14-targeted cancer cells. While a requirement for MMP14 is bypassed in low-density collagen hydrogels, invasion by the proteinase-deleted cells is associated with nuclear envelope and DNA damage, highlighting a proteolytic requirement for maintaining nuclear integrity. Nevertheless, when cancer cells confront explants of live human breast tissue, MMP14 is again required to support invasive activity. Corroborating these results, spatial transcriptomic and immunohistological analyses of invasive human breast cancers identified clear expression of MMP14 in invasive cells that were further associated with degraded collagen, underlining the pathophysiologic importance of this proteinase in directing invasive activity *in vivo*.

## Introduction

Invasion plasticity is a hallmark of cancer cells capable of assuming collective, mesenchymal or amoeboid phenotypes as they infiltrate host tissues (1–5). Mesenchymal-type phenotypes, elicited by leader cells during collective invasion or individual cells during single-cell invasion, assume elongated shapes, and exert contractile and protrusive forces on surrounding tissues (4, 6, 7). By contrast, amoeboid phenotypes are characterized by rounded tissue-invasive cells that move in a more fluid, Dictyostelium-like fashion as discrete single-cell units (1, 8, 9). Independent of the phenotype assumed, tissue-invasive cancer cells are confronted by the extracellular matrix (ECM), a heterogeneous composite of structural proteins, proteoglycans and glycosaminoglycans (10). Though complex, the ECM is dominated structurally by type I collagen, a triple-helical fibrillar protein that is deposited across a range of concentrations in a tissue-specific manner (11). Current dogma holds that cancer cells adopting a mesenchymal-type invasion program traverse dense ECM barriers by mobilizing proteolytic enzymes that actively degrade type I collagen rich-tissues (4, 12–14). By contrast, and perhaps in less restrictive environments, amoeboid cancer cells are thought to navigate similar tissues in a proteinase-independent fashion by using mechanical force to reversibly or irreversibly remodel the ECM, or alternatively, by distorting cell shape to a degree that allows them to negotiate otherwise size-restrictive pores (1, 4, 8, 15–18).

Here, we have used a combination of single-cell RNA sequencing, real-time imaging, 3-dimensional matrix models and human tissue explants to characterize the requirements of mesenchymal versus amoeboid cancer cell invasion in relation to collagen density, pore size and alignment as well as the modifying effects of nuclear rigidity, nutrient stress, hypoxia and cancer-associated fibroblasts. Unexpectedly, we find that while mesenchymal and amoeboid invasion mobilize distinct transcriptional programs, each activate a comparable set of ECM-degradative proteolytic enzymes. Further, cancer cells undergoing mesenchymal or amoeboid invasion both excavate tunnels through ECM barriers by proteolytically remodeling the surrounding type I collagen matrix in a pericellular fashion. In each case, a single membrane-anchored metalloproteinase, termed MMP14/MT1-MMP, plays a required role in driving tumor invasion. Following CRISPR/Cas9-mediated deletion of MMP14, cancer cells lose all tissue-invasive activity in dense matrix constructs while remaining unable to access an alternative, proteinase-independent invasive program. When confronting low-density collagen matrices, invasive activity proceeds independently of MMP14, but under these conditions, the proteinase nevertheless plays a critical, but previously unrecognized role in maintaining nuclear integrity and preventing DNA damage. Finally, when model collagen constructs are replaced with live human tissue explants, only MMP14-expressing cancer cells can actively tunnel through the surrounding matrix, a finding underscored in analyses of breast cancer spatial transcriptomics and tissue biopsies where invasive lesions are consistently found in association with MMP14 expressing-carcinoma cells decorated with tracks of type I collagen degradation products. Hence, both *in vitro* and *in vivo*, while invasion plasticity enables cancer cells to toggle between mesenchymal and amoeboid phenotypes, MMP14-dependent ECM proteolysis plays a critical role in driving tissue-invasive activity.

## Results

### Characterization of 3D mesenchymal versus amoeboid cancer cell invasion

Tumor spheroids comprised of human HT1080 fibrosarcoma cells embedded in 3D type I collagen hydrogels (2.2 mg/ml) with an average pore size of ∼2-4µm^2^ and rigidity of ∼162 Pa, actively infiltrate the surrounding matrix over a 72-hour culture period while displaying a largely collective/mesenchymal invasion program (Figure 1a-d, Movie 1). Characteristic of collagen-invasive, mesenchymal-like cells (6), control HT1080s demonstrate elongated cell shapes, associated with local strain on collagen fibers and activated β1-integrin (Figure 1e-i). As a decrease in β1-integrin activity induces amoeboid behavior, we cultured HT1080 spheroids in the presence of the β1-integrin blocking antibody, 4B4 (4, 12, 17). Under these conditions, 4B4-treated spheroids undergo a stark shift in behavior wherein features of collective/mesenchymal cell invasion are lost and replaced by invasive populations of single, rounded amoeboid cells (Figure 1a-i, Movie 2). Of note, in the presence of the blocking antibody, HT1080 cells maintain comparable invasive potential, while exhibiting characteristics of amoeboid activity, including increased levels of cortical actin, phospho-myosin light chain (pMLC) and plasma membrane blebbing (Figure 1j-n). Real-time cell-tracking analyses further support that mesenchymal and 4B4-treated amoeboid cells exhibit comparable average cellular speed and directional persistence (Figure 1 o-q, Movies 3 and 4).

**Figure 1:**
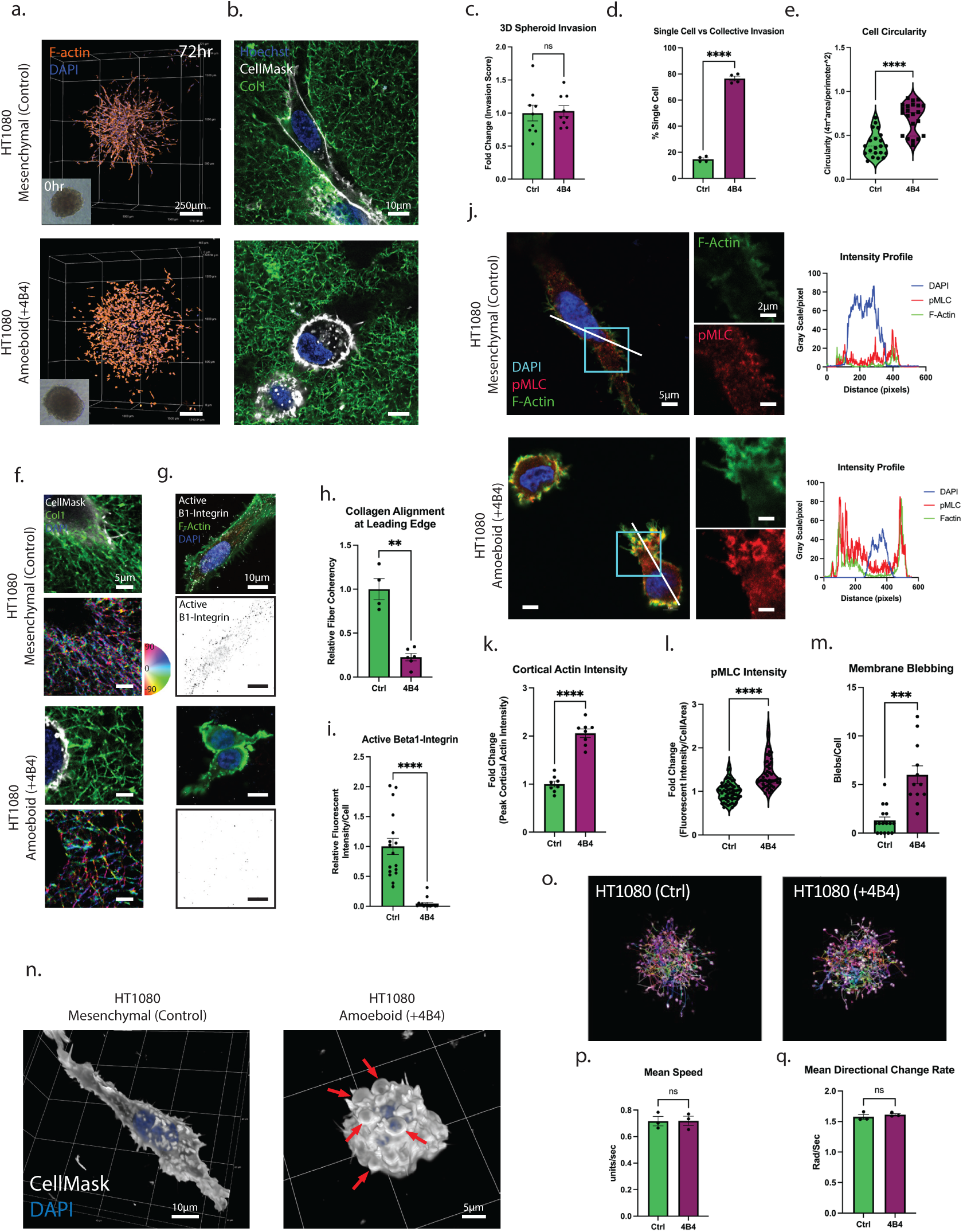
Characterization of Mesenchymal Versus Amoeboid Cancer Cell 3D Invasion. a) Representative three-dimensional fluorescent images of control and 4B4-treated (2µg/ml) HT1080 spheroids embedded in 2.2mg/ml type I collagen hydrogels after 72-hour culture and labelling with phalloidin and DAPI. Inset shows phase contrast images of spheroids at the time of embedding. b) High magnification images of control and 4B4-treated HT1080 cells invading through fluorescently labeled type I collagen hydrogels. Cells were labeled with CellMask-647 and Hoechst. c) Quantification of HT1080 invasion with and without 4B4-treatment. d) Quantification of the percentage of single vs collective invasive cells. e) Quantification of the circularity of individually invading tumor cells. f) Representative fluorescent images of collagen alignment observed at the leading edge of invasive tumor cells. Bottom images are colored based on fiber orientation. g) Representative fluorescent images of active ß1-integrin staining in invading tumor cells. h) Quantification of collagen alignment, by coherency of fluorescently labeled collagen fibers adjacent to invading cells. i) Quantification of detected ß1-integrin immunofluorescent staining. j) Immunofluorescent images of invading control and amoeboid cells labeled with phalloidin/DAPI and stained for pMLC. Intensity profiles, depict the fluorescent intensity by channel along the line designated in the image. k) Quantification of the peak cortical actin staining intensity in invading cells. l) Quantification of pMLC fluorescence in invading cells. m) Quantification of membrane blebs observed in single z-slice images of invading cells. n) Representative three-dimensional z-stack reconstructions of mesenchymal and amoeboid cells, labeled with Cellmask-647 and DAPI. Red arrows indicate membrane blebs on the cell surface. o) Endpoint images of invading HT1080 spheroids expressing GFP-NLS protein used to track nuclei during invasion. p,q) Quantification of mean speed and mean directional change rate detected during real time imaging of HT1080 invasion. P-value < 0.05 (*), <0.01 (**), <0.001 (***), < 0.0001 (****).

### Single-cell mapping of the mesenchymal and amoeboid invasion transcriptome

To begin characterizing the transcriptional programs that underlie 3D mesenchymal versus amoeboid invasion in an unbiased fashion, control and 4B4-treated spheroids were subjected to single-cell RNA sequencing after a 72-hour culture period. At this time point, cells are detected in a continuum of states, from stationary cells, still confined to the boundaries of the spheroid, to those most robustly migrating away from the 3D spheroid body (Figure 2a). Following cell cycle regression, initial clustering yielded three clusters for both mesenchymal and amoeboid conditions (Figure 2a). Consistent with the observed phenotypic progression of cells from stationary to highly motile states, visualization of gene expression associated with a cell migration gene set (19) demonstrated a gradual increase in migratory behavior (Figure 2a & Supplementary Figure 1a-c). These observations are further supported by additional analyses of Gene Ontology (GO) biological process pathways in merged control and 4B4-treated samples, where upregulated pathways associated with migration as well as tumor-ECM interactions concentrated in cluster 3, relative to clusters 1 and 2 (Supplementary Figure 1d). Upon comparing the identified motile populations of the mesenchymal and amoeboid conditions, 3227 genes are upregulated, and 917 genes down regulated in amoeboid vs mesenchymal-type invading cells (Figure 2b; top up/down DEGs). Top distinguishing genes by phenotype included *FOXD1*, *GAS6*, *STMN1* and *TENM2* in mesenchymal cells and *SPP1*, *ROMO1*, *ME1* and *FN1* in amoeboid cells (Figure 2c). Meanwhile, comparing conditions using a previously described GO amoeboid gene set (19) supported the amoeboid behavior of the 4B4-treated cells (Figure 2d). Key pathways described to contribute to an amoeboid phenotype, such as hypoxia-like HIF1-alpha signaling (4, 12, 20) and TGF-beta signaling (21), were likewise elevated in 4B4-treated cells (Figure 2d). Of note, amoeboid cells exhibited lower proliferative potential as demonstrated by fewer G2/M phase cells and decreased *PCNA* levels as well as lower cell numbers isolated following a 72-hour invasion period (Supplementary Figure 1e and f). Unexpectedly, upon interrogation of migration-associated genes, matrix metalloproteinases (MMPs), believed to play little, if any, role in amoeboid invasion, exhibited similar and even slightly elevated, expression levels in amoeboid-type invading cells relative to mesenchymal-type invading cells, including the membrane-anchored MMPs, *MMP14*, *MMP16*, *MMP17* and *MMP24* as well as the secreted MMP, *MMP2* (Figure 2d and e).

**Figure 2:**
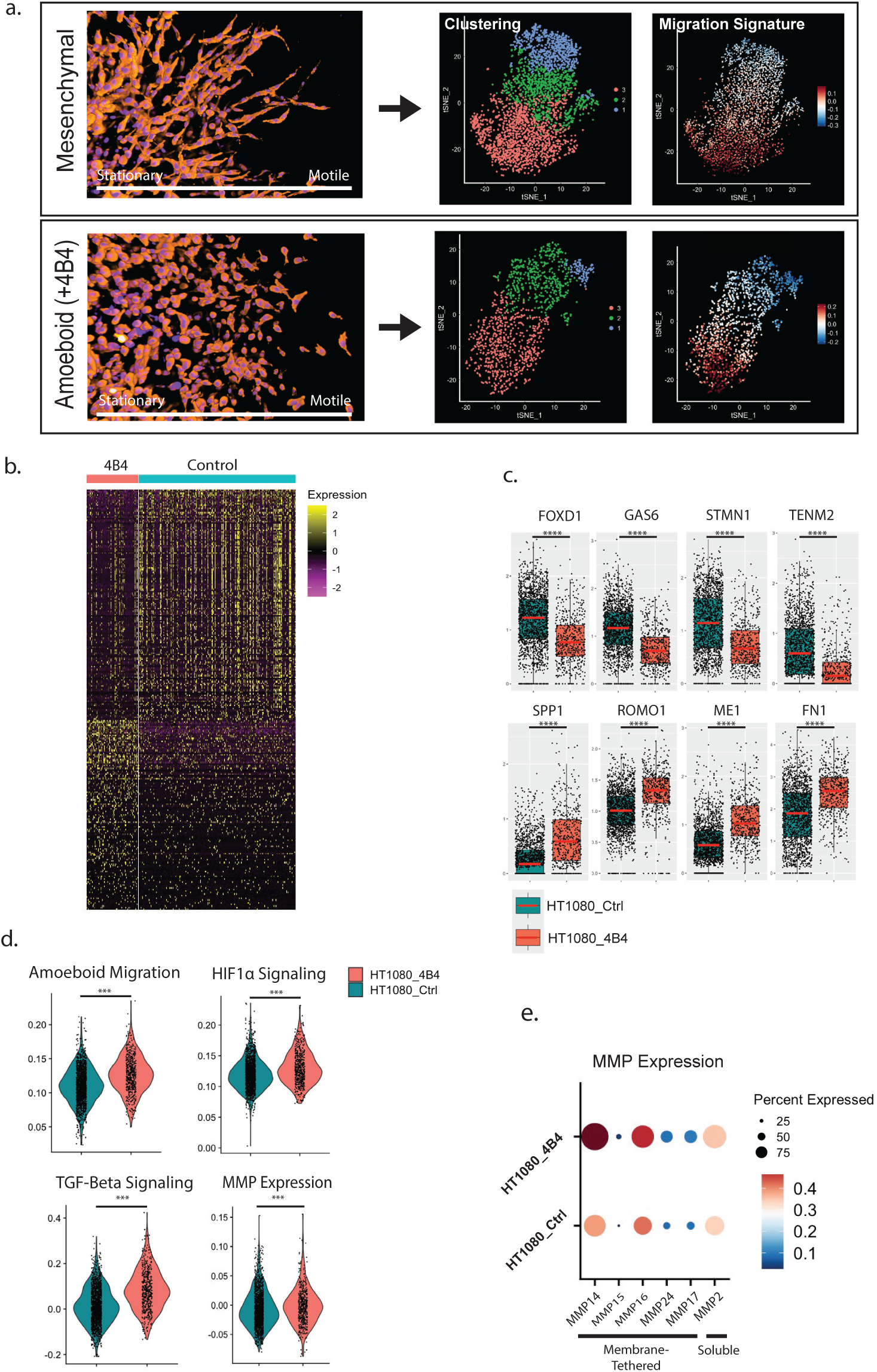
Single-Cell Transcriptomic Characterization of Mesenchymal and Amoeboid Invasion. a) Representative fluorescent images of HT1080 spheroids at 72 hours post-embedding, when single-cell RNA sequencing was performed. tSNE plots demonstrate initial clustering results and expression profiles of a cell migration gene set. b) Top differentially expressed genes between invading control cells and invading 4B4-treated cells. c) Box plots of top DEGs specific to each condition. Red lines indicate median values. d) Violin plots comparing gene expression of specified gene sets between control and 4B4-treated invading cells. e) Dotplot of MMP expression for control and 4B4-treated HT1080 cells. P-value < 0.05 (*), <0.01 (**), <0.001 (***), < 0.0001 (****).

### Mesenchymal- and amoeboid-type cell invasion requires MMP-dependent activity

Given that both mesenchymal and amoeboid invasion are associated with MMP expression in 3D culture, we sought to determine whether structural changes are occurring in the cross-linked collagen framework confronting the infiltrating cell populations. Following embedding of spheroids in fluorescently labelled 3D type I collagen hydrogels, matrix-free tunnels are detected, not only in the wake of invading mesenchymal cells, but amoeboid cells as well (Figure 3a and b). Moreover, as several MMPs are able to cleave collagen fibrils at a single site located approximately ¾ of the distance from its N-terminus (22), an antibody specific for this MMP-derived cleavage product, termed ¾ collagen, was utilized to detect MMP-dependent collagenolysis (23). Indeed, regardless of invasive strategy, invading cells were tightly associated with ¾-collagen fragments (Figure 3a and c), indicating that MMP-mediated proteolysis of type I collagen occurs during both mesenchymal and amoeboid invasion.

**Figure 3:**
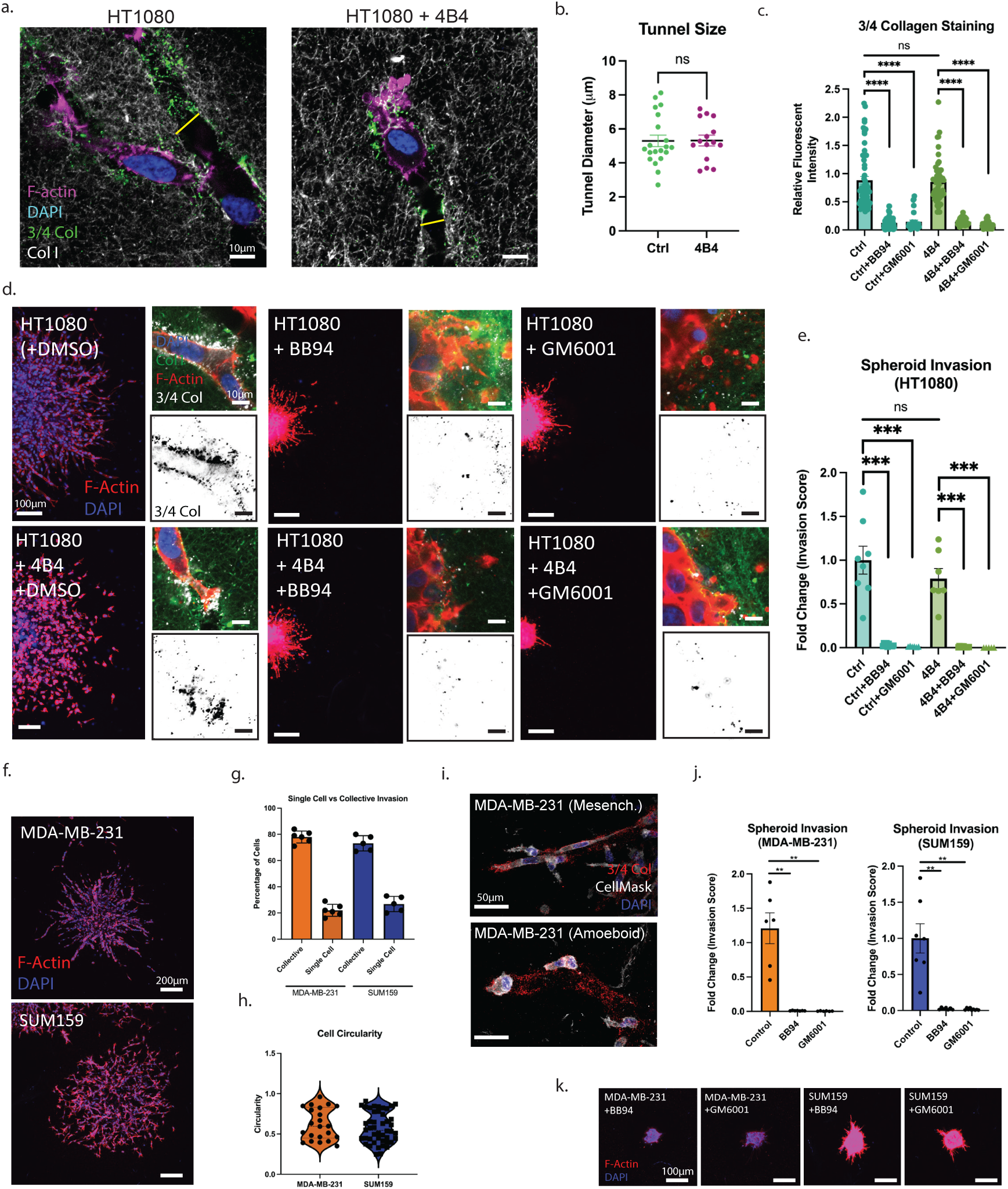
MMP-dependent Mesenchymal and Amoeboid Invasion. a) Representative immunofluorescent images of invading control and 4B4-treated HT1080 cells in fluorescently labeled collagen, stained for ¾ collagen degradation products. Yellow lines indicate tunnels formed in the collagen hydrogel. b, c) Quantification of tunnel diameter and fluorescent intensity of cell-associated ¾ collagen fluorescence. d) Fluorescent images of HT1080 spheroids after 72-hour culture in collagen in the presence of pan-specific MMP inhibitors BB-94 (5µM) or GM6001 (10µM). High magnification images, and inverted single channel images, show ¾ collagen detected at the cell-collagen interface. e) Quantification of spheroid invasion in the presence and absence of each MMP inhibitor. f) Representative images of invasive spheroids of MDA-MB-231 and SUM159 cells after 72-hours of invasion in collagen hydrogels. g,h) Quantification of single vs collectively invading cells and invasive cell circularity. i) Immunofluorescent images of spontaneously mesenchymal and amoeboid MDA-MB-231 cell invasion and associated ¾ collagen degradation products. j) Quantification of MDA-MB-231 and SUM159 spheroid invasion in the absence or presence of each MMP inhibitor. k) Fluorescent images of breast cancer spheroids treated with each MMP inhibitor. P-value < 0.05 (*), <0.01 (**), <0.001 (***), < 0.0001 (****).

To determine whether the observed collagenolytic activity is a necessary component of mesenchymal and amoeboid invasion, control or β1-integrin targeted HT-1080 spheroids were cultured in 3D hydrogels in the presence of either of two structurally distinct broad-spectrum MMP inhibitors, BB-94 or GM6001 (24). Contrary to reports of a ‘switch’ to proteinase-independent invasion in the face of proteinase inhibition (2, 8, 14, 15, 25), control HT1080 cells exhibited a complete block in invasion upon treatment with either MMP inhibitor (Figure 3d and e). While MMP-inhibited cells are able to extend thin filopodia-like protrusions and even release invasive cytoplasts, i.e., cytoplasmic fragments devoid of nuclei (26), into the surrounding matrix, movement of nuclei-containing, intact cell bodies into the surrounding matrix is inhibited completely (Figure 3d & e, Movies 5 & 6). Interestingly, 4B4-treated, amoeboid HT1080 cells are likewise unable to engage in proteinase-independent invasion in the presence of either BB-94 or GM6001 (Figure 3d and e). Loss of invasive potential in both cases is further associated with abrogated detection of ¾-collagen degradation products (Figure 3c and d). A similar complete block in invasion was observed using BB-94 in a ‘2.5D’ invasion assay, where cells are plated atop a 3D collagen hydrogel and allowed to invade into the subjacent matrix, a model often used to mimic the initiation of invasion as tumor cells breach the 2D-like barrier presented by the basement membrane (Supplementary Figure 2a and b) (27). To address whether MMP-inhibition is only sufficient to disrupt the initial migration away from the spheroids or if it would likewise halt the forward penetration of cells already committed to invasion, BB-94 was added after the spheroids were allowed to infiltrate the surrounding matrix for 48 hours. Following an additional 48-hour culture period in the presence of BB-94, invading cells halted in place, allowing lagging cells to ‘fill-in’ behind leading cells, but without progression of the leading edge (Supplementary Figure 3a-c). Thus, allowing embedded cells to stretch, exert cytoskeletal tension and activate mechanotransduction is not sufficient to maintain invasive activity in the absence of the continuous mobilization of MMP14 activity. Of note, even spheroids held in close proximity to one another, previously described to generate tension-induced alignment of collagen fibers and to allow for proteinase-independent invasion (28), remain susceptible to the delayed addition of MMP-inhibitors (Supplementary Figure 3a). Finally, as amoeboid activity has additionally been shown to be induced by nutrient stress (4, 12, 20), HT1080 spheroids in 3D culture were subjected to low glutamine, low-glucose or hypoxic conditions (Supplementary Figure 4a). Though varying degrees of a shift to single-cell amoeboid invasion are observed (Supplementary Figure 4a-d) under these conditions, cells remain reliant on MMPs as illustrated by the complete loss of invasive potential in the presence of BB-94 (Supplementary Figure 4a and b).

While β1-integrin blockade or nutrient stress allow for a transient induction of amoeboid behavior in HT1080 cells, we next sought to determine whether a similar reliance on MMP activity is observed in cancer cells capable of spontaneously engaging amoeboid phenotypes. In 3D spheroid culture, the human breast cancer cells lines, MDA-MB-231 and SUM159, likewise invade extensively into the surrounding type I collagen matrix, but with cells inherently exhibiting a mixture of mesenchymal and amoeboid phenotypes (29–31) (Figure 3f). In this system, ∼20% of MDA-MB-231 and ∼25% of SUM159 cells at the invasive front display amoeboid activity invading as single, rounded cells while the remainder migrate in a predominantly collective fashion (Figure 3f-h). Under these conditions, the collagen-invasive activity of both amoeboid and mesenchymal cells is associated with the generation of ¾-collagen degradation products (Figure 3i). Moreover, despite eliciting spontaneous amoeboid/collective phenotypes, invasion by both breast cancer cell lines is completely inhibited by BB-94 or GM6001 (Figure 3j and k, Movies 7 and 8).

### MMP14 directs both mesenchymal and amoeboid 3D invasion

As single-cell RNA sequencing indicated that mesenchymal and amoeboid invasion are associated with the expression of both secreted and membrane-anchored MMPs, we next sought to identify the MMP(s) responsible for ¾ collagen generation and invasion. To begin, we treated 3D-embedded spheroids with endogenous inhibitors of MMP activity, i.e., either TIMP-1 or TIMP-2 (32). TIMP-1, an effective inhibitor of secreted MMPs (32), did not affect invasion while TIMP-2, an inhibitor of both secreted and membrane-tethered MMPs (32), strongly blocks invasion of HT1080, MDA-MB-231 or SUM159 cells (Supplementary Figure 5a-c). Of the MMPs detected, the membrane-tethered MMP, *MMP14/*MT1-MMP, is known to function as an effective pericellular type I collagenase (33–36). To monitor endogenous MMP14 production and trafficking in real-time, we generated a knock-in *MMP14* allele with an mCherry sequence inserted into the extracellular juxtamembrane domain of the proteinase (Figure 4a). Following the generation of the MMP14^mCherry-KI^ HT1080 and MDA-MB-231 lines, live cells cultured atop collagen hydrogels actively trafficked MMP14^mCherry-KI^ containing vesicles throughout the cell body (Figure 4b). To differentiate cell surface-associated vs intracellular MMP14, cells were immunostained using an mCherry-specific antibody before or after cell permeabilization. In both permeabilized mesenchymal and amoeboid cells, perinuclear accumulation of MMP14^mCherry-KI^ is observed with substantial amounts of MMP14^mCherry-KI^ additionally found in vesicles distributed throughout the cytoplasm as well as the cell periphery (Figure 4c and Supplementary Figure 6). In non-permeabilized cells, MMP14^mCherry-KI^ exclusively outlines the cell surface (Figure 4c and Supplementary Figure 6) and when visualized as a 3D-reconstruction, highlights the surface presentation of the proteinase in both mesenchymal and amoeboid cells (Figure 4d).

**Figure 4:**
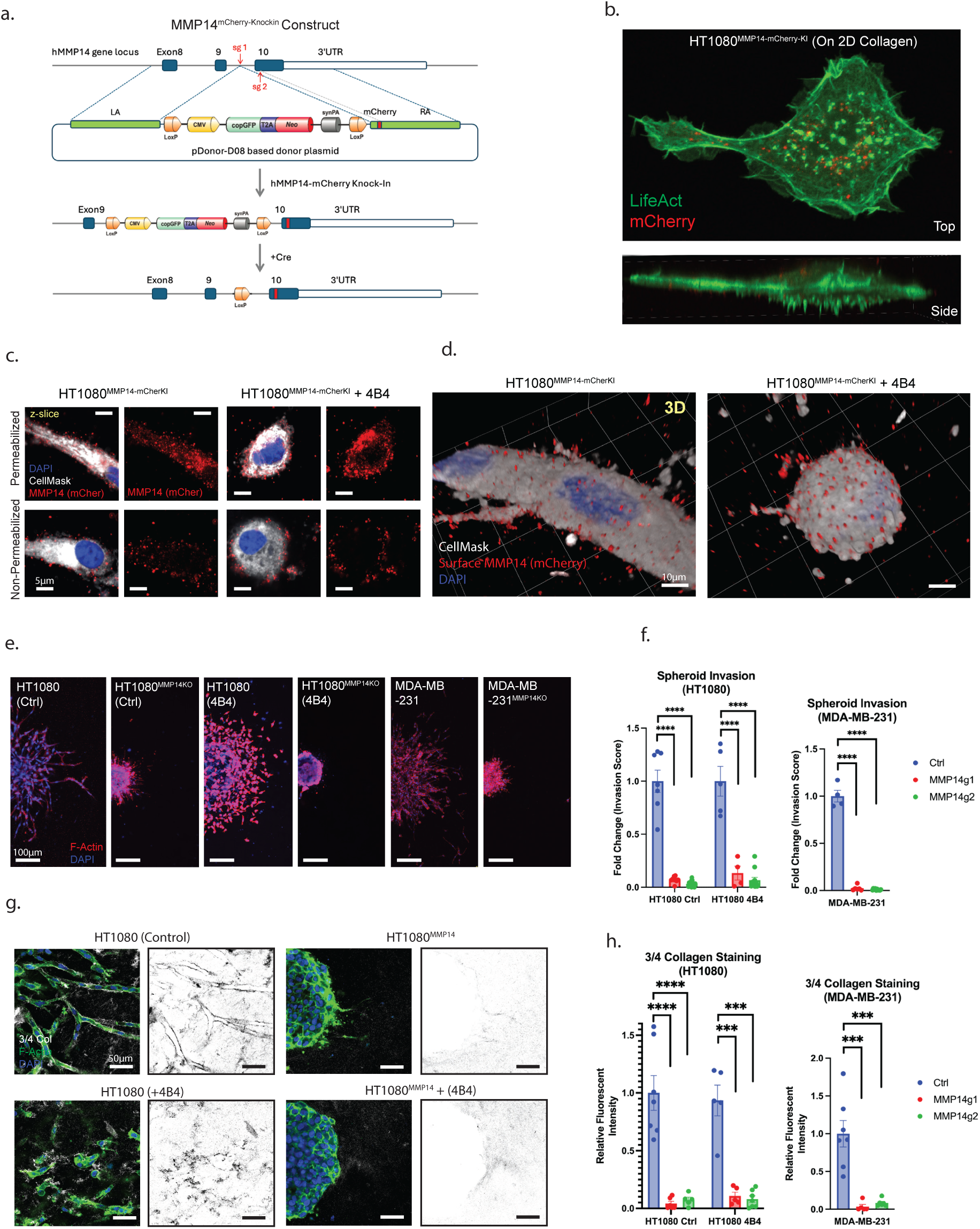
MMP14 Directs Mesenchymal and Amoeboid 3D Invasion. a) Depiction of the constructs used to generate MMP14-mCherry knock-in HT1080 and MDA-MB-231 cells. b) Representative 3D images of an HT1080^MMP14-mCherryKI^ cell transfected with GFP-LifeAct atop a thin collagen gel. c) Immunofluorescent images of permeabilized and non-permeabilized HT1080 cells stained with an mCherry-specific antibody to label total vs cell surface MMP14-mCherry proteins. d) 3D reconstructions of z-stack images of non-permeabilized control and 4B4-treated HT1080 cells stained for mCherry. e) Representative images of control and MMP14-KO spheroids after 72 hours of invasion. f) Quantification of spheroid invasion with CRISPR targeting of MMP14 via two separate guide RNAs. g) Immunofluorescence staining of ¾ collagen associated with control and MMP14-targeted spheroids. h) Quantification of ¾ collagen staining at the tumor cell-collagen interface. P-value < 0.05 (*), <0.01 (**), <0.001 (***), < 0.0001 (****).

With clear surface localization of MMP14 observed in invading mesenchymal and amoeboid cells, we assessed its impact on each invasive program, using CRISPR/Cas9 to delete its expression in HT-1080, MDA-MB-231 and SUM159 cells (Supplementary Figure 7a). Following targeting with two distinct guide RNAs, all MMP14-deleted cell lines retain normal proliferative and motile activity in standard 2D culture (Supplementary Figure 7b and c). By contrast, spheroids prepared from knockout HT-1080 cells – either in the absence or presence of β1-integrin targeting – are devoid of 3D collagen-invasive activity while failing to generate ¾ collagen degradation products despite inserting filopodia-like protrusions into the surrounding matrix, (Figure 4e-h, Movie 9). Similar, if not identical, results are observed with MMP14-targeted MDA-MB-231 or SUM159 spheroids (Figure 4e-h, Supplementary Figure 7d, Movies 10), thereby highlighting its absolute requirement for driving invasive activity. To rule out the possibility that an MMP14-derived cleavage product yields a pro-motility signal that is lacking in MMP14-null spheroids, or alternatively, that MMP14-null cells generate an inhibitory signal that would impede wild-type cancer cell invasion, control and MMP14-null tumor spheroids were assembled separately and then cultured in close proximity within 3D collagen hydrogels in order to expose each population to soluble factors generated by the opposing population. However, co-culture neither rescues the invasive potential of MMP14-null cells nor interferes with the activity of control spheroids that continue to invade extensively (Supplementary Figure 8a). Only when control and MMP14-null cancer cells are co-mixed into single spheroids were MMP14-null cells observed invading in 3D, presumably by following control cells acting as leader-cells (Supplementary Figure 8b). Taken together, these results are most consistent with a parsimonious model wherein MMP14 functions as a *cis*-acting proteinase that drives invasive activity by cleaving barrier-forming ECM components directly confronted by the cell. Consistent with this conclusion, wild-type MMP14-expressing cancer cells are unable to infiltrate synthetic PEG hydrogels unless crosslinks incorporate MT1-MMP scissile bonds (37) (Supplementary Figure 9a and b) while wild-type cell invasion into synthetic hydrogels incorporating cleavable moieties is blocked in the presence of pan-specific MMP inhibitors or when MMP14 is specifically deleted (Supplementary Figure 9a and b).

### Collagen cross-linking rather than pore size dictates proteinase-independent invasive potential

While we find that cancer cells confronting pore sizes on the order of ∼2-4um^2^ in native collagen hydrogels are unable to mount invasive behavior by MMP14-independent mechanisms, a body of work suggests that proteinases are no longer required to support invasion when average pore sizes exceed that of ∼10% of the cell’s nuclear dimensions, i.e., roughly 7um^2^ in HT-1080 and MDA-MB-231 cells (5, 14). Pepsinized collagen (atelocollagen), lacking the non-helical telopeptide domains critical for intermolecular crosslinking between collagen fibers, has frequently been used to generate larger pore constructs conducive to the study of proteinase-independent invasion *in vitro* (8, 12, 20, 21, 38–40). Indeed, atelocollagen-based constructs allow for the formation of high-porosity hydrogels with an average pore size of ∼17um^2^, well in excess of the proposed nuclear size restrictions of HT-1080 or MDA-MB-231 cells (14) (Figure 5a and b). Under these conditions, both MMP14-wild-type and MMP14-null HT-1080 and MDA-MB-231 spheroids exhibit strong invasive potential (Figure 5c-d). Interestingly, despite forgoing the requirement for matrix remodeling in atelocollagen hydrogels to support invasion (Figure 5c-d), ¾ collagen degradation products are still detected as wild-type cells invade (Supplementary Figure 10). By contrast, the invasive activity of MMP14-null cancer cells proceeds in the absence of ¾ collagen degradation products (Supplementary Figure 10). While these observations of proteinase-independent invasion of atelocollagen concur with previous reports (8, 13–15), it remains unclear as to whether proteinase-independent invasion arises as a consequence of the increase in matrix pore size, or the absence of the covalent collagen cross-links characteristic of native tissues (41). As such, we alternatively generated telopeptide-intact type I collagen hydrogels formed at low temperatures to slow fibrillogenesis, thereby allowing for the generation of large pore constructs comparable to those observed in atelocollagen matrices (Figure 5a and b) (14, 42, 43). Under these conditions, while both control HT-1080 and MDA-MB-231 spheroids invade robustly in the high porosity native collagen gel, MMP14-null cancer cells display almost no invasive potential despite the larger pore size, aside from small numbers of cells that only partially separate from the spheroid but remain in close proximity to the central cell mass (Figure 5c-d). Hence, even in type I collagen matrices with pore sizes exceeding those previously held to be permissible to proteinase-independent invasion, mesenchymal as well as amoeboid activity remains MMP14-dependent when the covalent cross-links that define native interstitial matrices are present.

**Figure 5:**
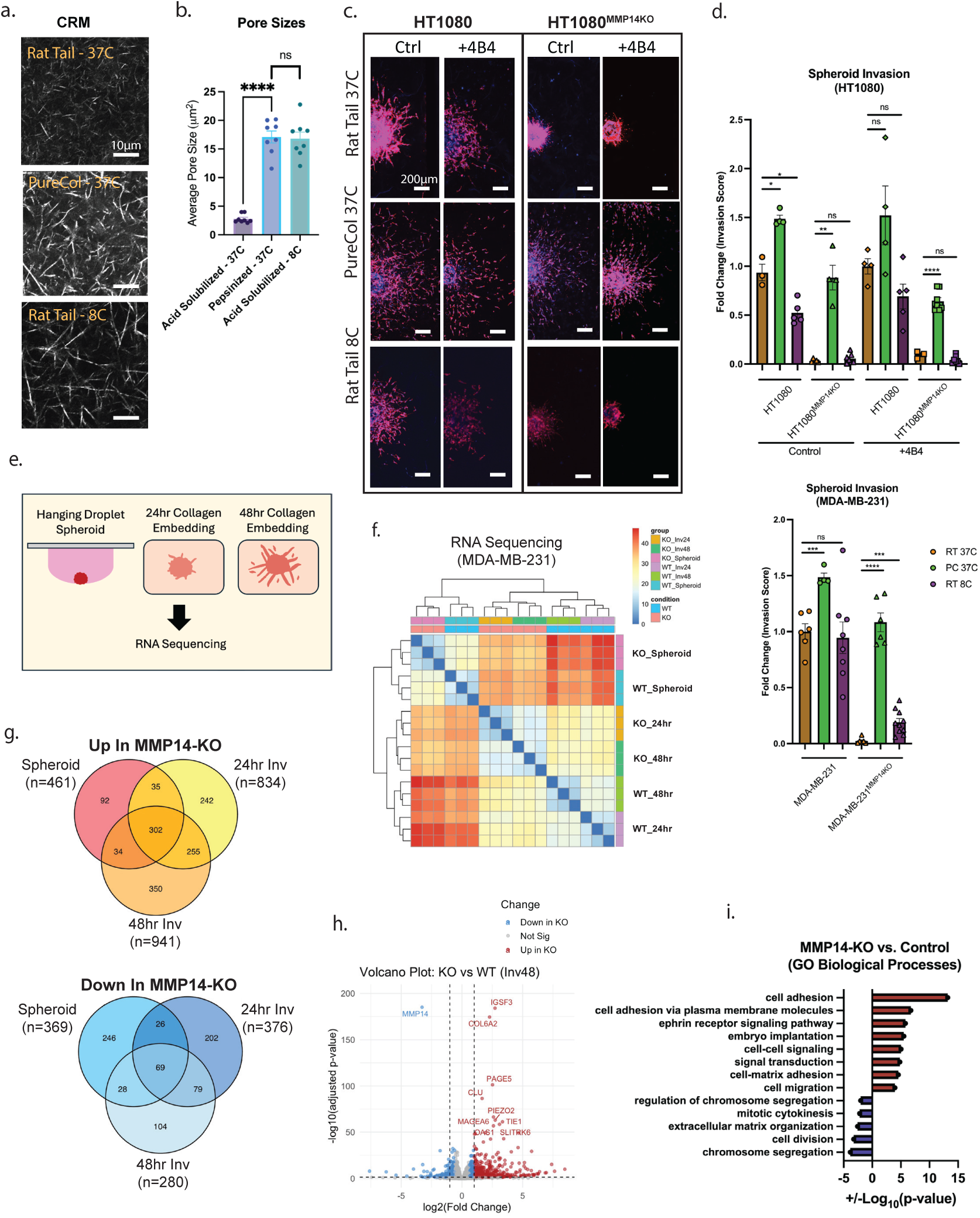
MMP14-dependent Cancer Cell-Collagen Interactions and Transcriptional Regulation. a) Confocal reflectance microscopy images of collagen hydrogels prepared using acid-solubilized rat collagen gelled at 37°C or 8°C and pepsin-extracted bovine collagen (PureCol) gelled at 37°C. All hydrogels contain 2.2mgl/ml collagen. b) Quantification of average collagen pore sizes in each condition. c) Representative fluorescent images of invading spheroids in the indicated collagen conditions. d) Quantification of spheroid invasion in the indicated collagen conditions. e) Illustration of the three time points included in RNA sequencing of control and MMP14-KO MDA-MB-231 spheroids. f) RNA sequencing sample association heat map, grouped by transcriptional similarity. g) Venn Diagrams of genes up and down regulated in MMP14-targeted MDA-MB-231 spheroids versus controls at each timepoint. h) Volcano plot of differentially expressed genes between MMP14-KO and control spheroids after 48 hours in collagen. i) Gene Ontology biological processes up and down regulated in MMP14-KO spheroids compared to controls after 48-hour culture in collagen. P-value < 0.05 (*), <0.01 (**), <0.001 (***), < 0.0001 (****).

While the loss of intact collagen cross-linking rather than pore size is the key matrix element allowing for proteinase-independent invasion in collagen I hydrogels, these results emphasize the ability of MMP14-null cells to effectively mount 3D invasive activity under permissive conditions. However, the impact of blocked invasion observed in MMP14-null cells on transcriptional programs remains undefined. As such, high-throughput RNA sequencing was performed on wild-type vs MMP14-targeted MDA-MB-231 spheroids upon initial spheroid formation as well as following 24 and 48 hours embedded in intact type I collagen hydrogels (Figure 5e and f). Rather than becoming progressively more quiescent, immobile MMP14-null cells up regulate transcript levels of a large family of genes relative to invading controls over the course of the experiment (Figure 5g). Focusing on the differences observed following 48 hours in collagen, while genes associated with proliferation are in fact downregulated in MMP14-null cells, transcripts related to cell-cell adhesion, cell-ECM adhesion and cell migration are all elevated in the MMP14-KO cells, suggesting an enhanced effort of targeted cells to mount an invasive response while being limited by their inability to cleave surrounding collagen molecules (Figure 5h-i). Of note, comparing all spheroid conditions, expression of other MMPs remain largely consistent in MMP14-null cells, while *MMP2*, *MMP9* and *MMP10* levels increase in MMP14-null cells (Supplementary Figure 11).

### Cancer cell- and microenvironment-intrinsic modulation of 3D invasion programs

Independent of MMP14 activity, recent reports suggest that changes in either cancer cell phenotype or the surrounding tumor microenvironment can modulate the matrix remodeling requirements necessary for 3D invasion. With regard to cancer cell-intrinsic mechanisms, the nucleus serves as the most rigid intracellular organelle whose mechanical properties dictate the ability of motile cells to traverse limiting pore dimensions (14, 18, 44). As nuclear rigidity and deformability are largely controlled by the nuclear lamina proteins, lamin A and C (products of the *LMNA* gene), we sought to determine if *LMNA-*deleted cells acquire proteinase-independent invasive activity (14, 16, 45–47). Following *LMNA* targeting in either HT-1080 or MDA-MB231 cells (Supplementary Figure 12a), cells transmigrate fixed 8.0 µm filter pores comparably to controls, but when confronted with subnuclear 3.0 µm pores, *LMNA*-targeted cancer cells demonstrate enhanced transmigration potential (Supplementary Figure 12b). However, unlike fixed pores constructed of synthetic material, pores found in collagen hydrogels are not aligned in register, thereby forcing migrating cells to negotiate a more tortuous path, encountering multiple, uniquely sized and shaped pores simultaneously. Indeed, in 3D collagen culture, while *LMNA* knockout HT-1080 spheroids display invasive activity comparable to controls in the presence or absence of β1 integrin targeting, invasion is completely blocked following MMP inhibition with comparable results obtained using *LMNA*-targeted MDA-MB-231 spheroids (Supplementary Figure 12c and d).

Independent of cancer cell-intrinsic changes in phenotype, alterations in chemotactic signals, matrix composition, fluid viscosity or alignment of collagen fibrils can modulate migratory activity (48–58). While EGF and SDF1 can serve as potent cancer cell chemoattractants (48, 49), neither stimulus rescues invasive activity in MMP14-null cancer cells (Supplementary Figure 13a and b). Type I collagen hydrogels impregnated with exogenous fibronectin (50, 51) or collagen VI (52) have each been reported to enhance 3D cancer cell invasion programs with similar activity assigned to increases in fluid viscosity (30). However, under each of these conditions, MMP14-targeted cells remain unable to infiltrate the surrounding hydrogel (Supplementary Figure 13a and b). Of note, the physical alignment of collagen fibrils, particularly when tension-induced, has been reported to be conducive for proteinase-independent invasion (28, 53, 54). As such, we used a mechanical tension-inducing apparatus to allow for the controlled induction of focal collagen alignment (Figure 6a). When control and MMP14-targeted spheroids were cultured together in the central region of the aligned gels, control cancer cell spheroids preferentially invade parallel with the aligned fibers, while MMP14-targeted cells remain unable to invade despite the surrounding tension-aligned collagen (Figure 6b and c). Finally, in the absence of matrix-degrading proteolytic systems, invasion-incompetent cancer cells have been reported to interact with accessory cell populations, particularly cancer-associated fibroblasts (CAFs) (50, 55–58) which alternatively remodel the ECM via either mechanical or proteolytic processes, thereby allowing cancer cells to passively infiltrate surrounding tissues in *trans*. To this end, wild-type or MMP14 knockout MDA-MB-231 spheroids were embedded in collagen hydrogels containing patient-derived human breast CAFs (59). Consistent with their reported invasion-promoting properties, co-embedding wild-type spheroids with CAFs significantly enhances the invasive capacity of control cancer cells (Figure 6d and f). However, upon co-culture with MMP14-null MDA-MB-231 spheroids, CAFs are incapable of rescuing cancer cell invasion despite the fibroblasts’ ability to locally proteolyze surrounding collagen fibers (Figure 6d-g). Together, these results underline an inescapable reliance on cancer cell MMP14 to drive invasive activity through native collagen barriers.

**Figure 6:**
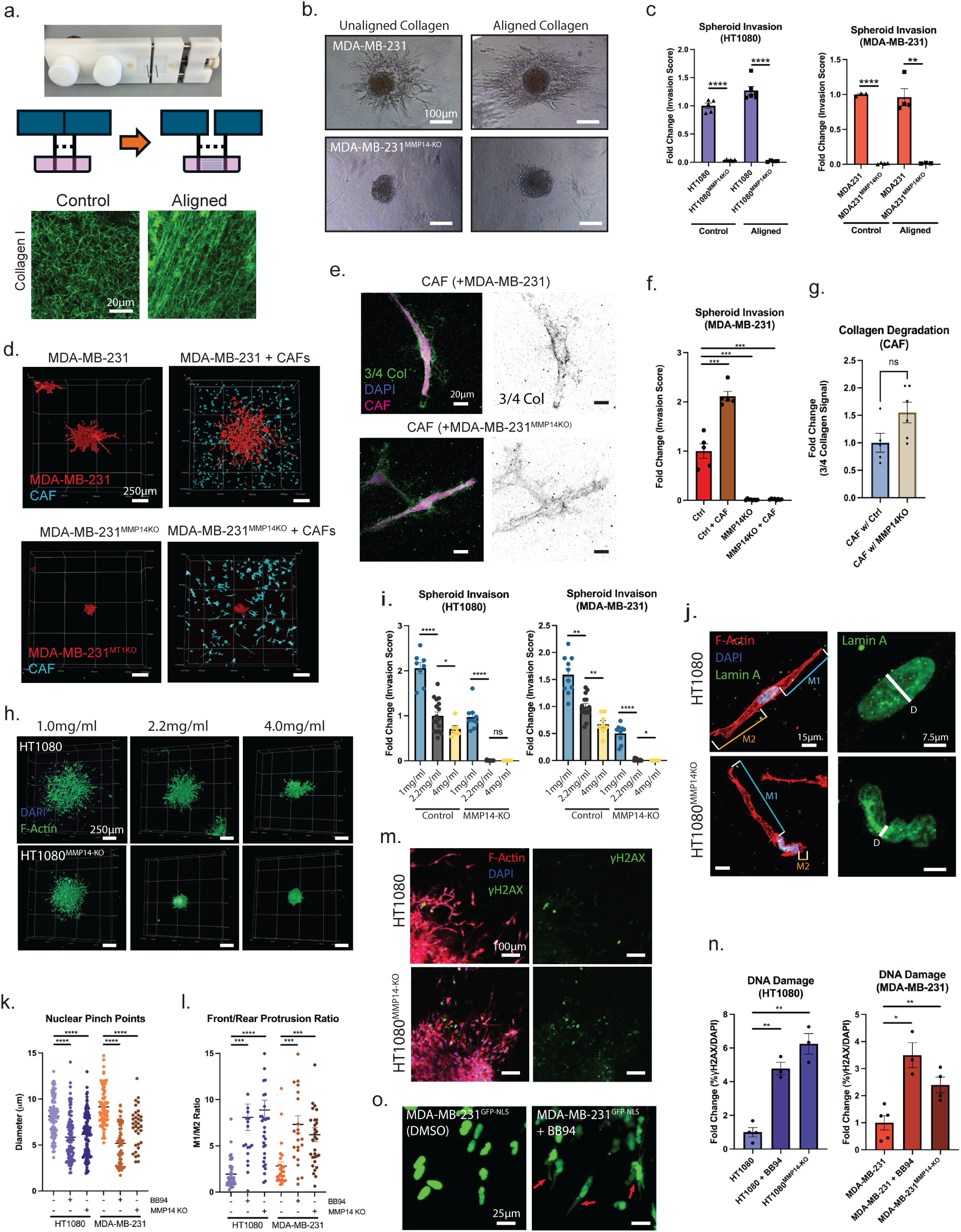
Regulation of MMP14-dependent Invasion by Cell- and ECM-Intrinsic Factors. a) Image and illustration of the apparatus used to mechanically apply tension to collagen hydrogels and align collagen fibers along with images of fluorescently labeled fibers with and without alignment. b) Phase contrast images of MDA-MB-231 spheroids in unaligned versus aligned regions of the collagen hydrogel. c) Quantification of control and MMP14-KO tumor spheroid invasion in aligned vs unaligned regions of collagen hydrogels. d) Three-dimensional images of MDA-MB-231 tumor spheroids following 72 hours of culture in the presence or absence of human breast cancer-associated fibroblasts (5X10^5^ cells/ml collagen). e) Immunofluorescent images of breast cancer CAFs cultured with control or MMP14-targeted MDA-MB-231 spheroids and stained for ¾ collagen degradation products. f,g) Quantification of MDA-MB-231 spheroid invasion in the absence or presence of CAFs along with the detection of collagen degradation associated with CAFs in each condition. h) Three-dimensional images of spheroid invasion in the indicated concentration of acid-solubilized rat tail collagen. i) Quantification of spheroid invasion in the indicated collagen densities. j) Representative immunofluorescent images of control and MMP14-KO HT1080 cells invading through 1mg/ml collagen. White line depicts measurements for nuclear pinch points. Indicated M1 and M2 measurements depict front and rear protrusion measurements used for quantification. k,l) Quantification of nuclear pinch points and front/rear protrusion ratio for the indicated cells conditions invading through 1mg/ml acid-solubilized rat tail collagen hydrogels. m) Representative immunofluorescent staining of γH2AX DNA damage sites. n) Quantification of γH2AX staining in cells invading through 1mg/ml collagen hydrogels. o) Representative fluorescent images of control and MMP14-targeted MDA-MB-231 cells expressing a GFP-NLS reporter invading through 1mg/ml collagen, red arrows indicate cells with nuclear envelope rupture. P-value < 0.05 (*), <0.01 (**), <0.001 (***), < 0.0001 (****).

### MMP14-dependent maintenance of nuclear integrity

*In vivo*, cancer cells may encounter collagenous barriers across a range of densities with increases in collagen concentrations and rigidity proposed to trigger the generation of more aggressive tumor phenotypes (60–62). As such, cancer cell spheroids were embedded in collagen hydrogels at 1.0 mg/ml, 2.2 mg/ml and 4.0 mg/ml where matrix rigidity and pore size range from ∼20Pa to ∼400Pa, and from ∼1µm^2^ to ∼30 µm^2^, respectively (Supplementary Figure 14a and b). Despite reports that increases in matrix rigidity induce cancer cells to assume more aggressive phenotypes, the invasive activity of HT-1080 and MDA-MB-231 spheroids decrease as a function of collagen concentration (Figure 6h and i). By contrast, the dominant collective cell invasion program observed in 2.2-4.0 mg/ml collagen gels shifts to a primarily single-cell pattern of invasion at 1.0 mg/ml collagen, supporting the emergence of cell-cell jamming when more restrictive matrix barriers are confronted (Figure 6h and Supplementary Figure 14c) (13, 63). Interestingly, while invasion of MMP14-null cancer cells is completely abrogated in higher density collagen gels, significant invasive activity is retained in low-density collagen matrices (Figure 6h and i), a finding supportive of earlier reports describing proteinase-independent invasive activity when collagen concentration is lowered in cross-linked hydrogels (64). Interestingly, despite the apparently dispensable role for MMP14 under these conditions, collagen-infiltrating cancer cells continue to actively degrade pericellular collagen in low-density gels (Supplementary Figure 15). Furthermore, while invading wild-type cancer cells display normal nuclear shapes in either low- or high-density collagen gels, MMP14-null tumor cells invade the low-density matrices in tandem with marked distortions in nuclear shape and position, as indicated by nuclear pinch points and the ratio of front and rear protrusion lengths, coincident with attendant increases in DNA damage as assessed by γH2AX staining (Figure 6j-n). Of note, neither control, nor MMP14-null cancer cells display similar changes in 2D culture (Supplementary Figure 16a). These observations suggest that in the absence of proteolytic activity, nuclei undergo significant shape changes to support invasion even in large pore, low-density collagen matrices. Consistent with this premise, the increased nuclear deformation observed in BB94-treated or MMP14-null invading cells is strongly associated with increased nuclear envelope rupture, visualized by bursts of GFP-NLS fluorescent signals appearing in the cytoplasmic compartment (Figure 6o and Movies 11 and 12). Hence, while MMP14 is not required to support invasion per se in low-density collagen hydrogels, the collagenolytic widening of matrix pores is required in order to maintain nuclear integrity.

### Proteinase requirements at the cancer cell-stromal interface of human tissue

Our findings demonstrate that cancer cells, though capable of moving between collective, mesenchymal and amoeboid phenotypes, are solely dependent on MMP14 to drive invasive activity in higher density collagen hydrogels and only demonstrate proteinase-independent invasion programs in low-density collagen matrices. However, native, reconstituted type I collagen hydrogels cannot fully recapitulate the more complex composition and architecture of the interstitial matrix. Therefore, we utilized live, patient-derived breast tissue explants to assess the relative potential of MMP14-wild-type cells to infiltrate native stromal barriers in the absence or presence of BB-94 as well as after MMP14 targeting. To avoid creating a damaged passageway through the tissue at an injection site, we used a 2.5D invasion format for these studies wherein culture of fluorescently labeled MDA-MB-231 cells atop the stromal surface led to frank invasion over a 6-day culture period (Figure 7a-c). In marked contrast, invasion is completely blocked when MMP activity is inhibited with BB-94 or when MMP14 is deleted (Figure 7b and c). Hence, when substituting human tissue for type I collagen hydrogels, a requirement for MT1-MMP activity in driving invasive activity is retained.

**Figure 7:**
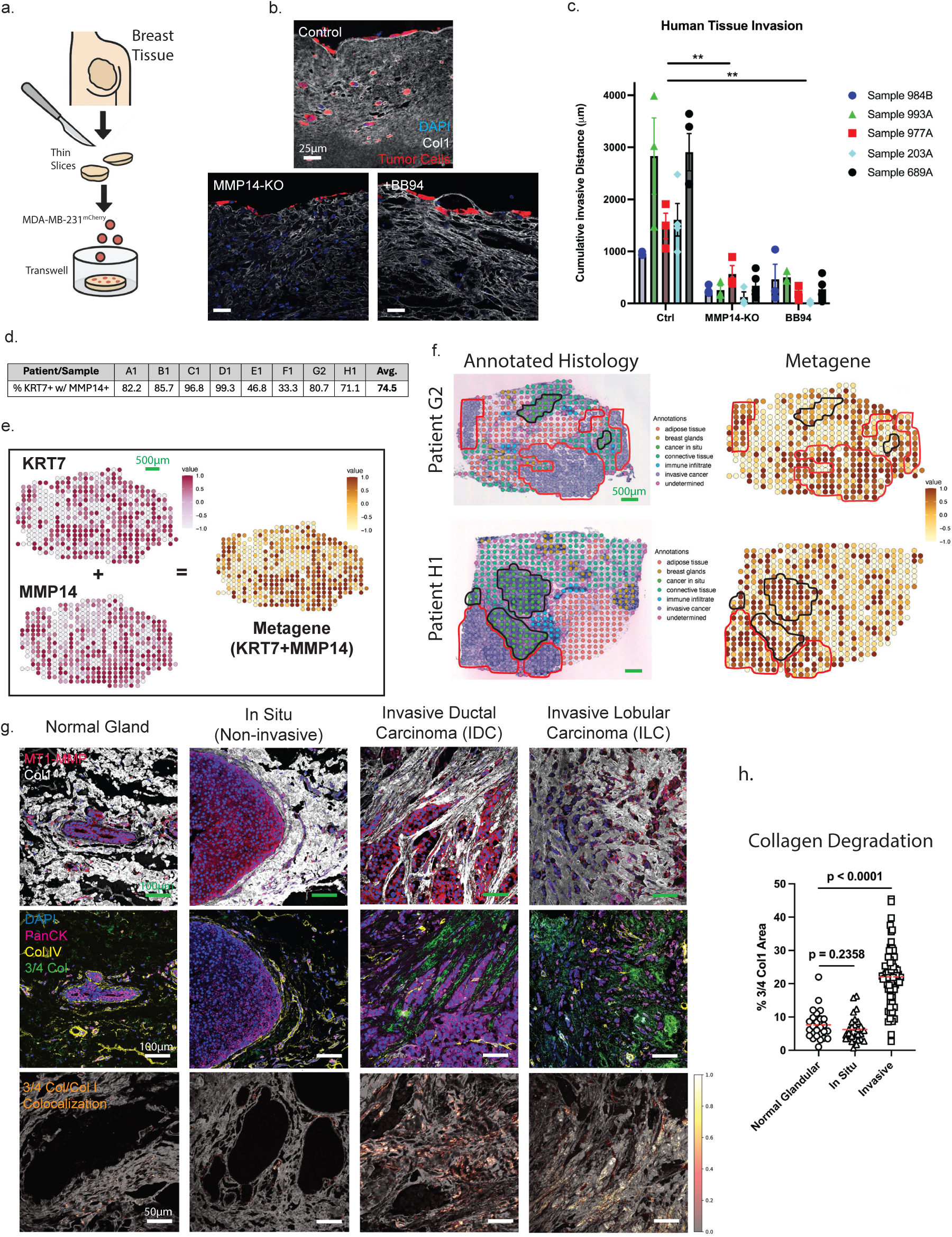
Proteinase Requirements at the Cancer Cell-Stromal Interface of Human Breast Tissue. a) Cartoon depicting experimental setup for *ex vivo* human breast tissue invasion assays. b) Representative immunofluorescence images of cross-sections of human breast tissue samples following 6-days of MDA-MB-231 cancer cell invasion. c) Quantification of tumor cell invasion into human breast tissue, grouped by individual patient sample (n=5). Average values for each sample were compared between conditions using a paired t-test. d) Table summarizing the percentages of KRT7-positive spots also positive for MMP14, based on normalized gene expression, within invasive cancer regions in a spatial transcriptomic dataset from 8 patient samples. e) Spatial plots depicting an example of KRT7 and MMP14 z-scores compared to the z-scores for the combined KRT7/MMP14 metagene. f) Two representative examples of annotated tissue sections compared to the corresponding KRT7/MMP14 metagene z-score plot. Red outlines depict invasive cancer regions and black outlines depict *in situ* cancer lesions. g) Representative immunofluorescent images of normal and malignant human breast tissues, stained for the indicated antibodies. The bottom row depicts sites of colocalization between type I collagen staining and ¾ collagen staining. h) Quantification of collagen degradation detected in human tissues. Measurements taken from 13 clinical samples with ∼10 regions assessed in each sample. P-value < 0.05 (*), <0.01 (**), <0.001 (***), < 0.0001 (****).

As our *ex vivo* data support a requirement for MMP14 for tumor cell invasion through human tissue-derived ECM, we next sought to assess clinical human cancer samples for the presence of *MMP14* expression. Indeed, using publicly available bulk RNA sequencing data comparing 2881 normal and 4484 tumor samples from common solid tumor types (65), including bladder, breast, colon, liver, lung (small cell), pancreas, renal clear cell and papillary types, skin, stomach and testis, *MMP14* is consistently detected across the various tissue types. Furthermore, *MMP14* is significantly elevated in tumor tissues (Supplementary Figure 17a and b) While these data support the potential clinical relevance of MMP14 across a range of cancers, bulk sequencing does not specifically evaluate tumor cell MMP14 expression or activity. As such, we sought to evaluate MMP14 mRNA and protein expression levels, specifically in invasive breast cancer lesions. Using publicly available spatial transcriptomic data from breast cancer patients (66), we find that the majority of sites positive for the epithelial marker, *KRT7*, are likewise *MMP14*-positive (>70%) (Figure 7d). A metagene identifying *MMP14*-positive epithelial cells (*MMP14*+/*KRT7*+) was used to map areas predicted to contain invasive regions in an unbiased fashion (Figure 7e). Indeed, regions annotated based on histology as invasive cancer, consistently displayed high *MMP14*/*KRT7* metagene z-scores (Figure 7f and Supplementary Figure 18, invasive regions outlined in red), supporting *MMP14* expression in invasive breast cancer cells. Of note, varying levels of the *MMP14*/*KRT7* metagene were also detected in *in situ* cancer regions present in 3 of the 8 patient samples (Figure 7f and Supplementary Figure 18, *in situ* regions outlined in black).

Finally, as spatial transcriptomic data is nevertheless limited to detection of *MMP14* mRNA, we further aimed to determine whether the joint appearance of MMP14 protein and degraded type I collagen could be found at sites of invading breast cancer cells in clinical samples. In comparisons between normal breast tissue, *in situ*/non-invasive tumor regions and frankly invasive lesions, we detect MMP14 in both normal and malignant mammary epithelial cells (Figure 7g). However, upon staining of the basement membrane protein, type IV collagen, we confirm continuous basement membrane surrounding both normal and non-invasive malignant regions (67), suggesting that epithelial cells in these areas do not establish direct contact with the underlying interstitial matrix (Figure 7g). However, in areas devoid of basement membrane staining that signal the emergence of invasive carcinomas, MMP14-positive cancer cells are found in association with large deposits of ¾ collagen degradation products (Figure 7g and h). By contrast, in normal breast tissue or non-invasive carcinomas surrounded by an intact basement membrane, type I collagen degradation fragments are seldom observed as confirmed by quantitative analysis (Figure 7g and h). Interestingly, as invasive lobular carcinomas (ILCs) have been reported to display more amoeboid-like, single-cell invasion programs (68, 69) as opposed to the predominantly collective/mesenchymal invasion observed in invasive ductal carcinomas (IDCs) (13), we specifically assessed whether MMP14 and ¾ collagen were detected in both subtypes of invasive breast cancer. Indeed, invasion-associated MMP14 and ¾ collagen degradation products are readily detected in both IDC (n=9) and ILC (n=3) (Figure 7g). Taken together, these findings support a model wherein cancer cells invade type I collagen-rich tissues *in vitro* and *in vivo* by mobilizing the membrane-anchored metalloproteinase, MMP14, to proteolytically remodel the pericellular matrix to accommodate cell trafficking.

## Discussion

Invasion plasticity imbues cancer cells with the ability to penetrate tissue barriers by reversibly adopting any one of a range of distinct motility programs (1, 4, 70). Current dogma holds that collective as well as single-cell patterns of invasion can be supported by either mobilizing matrix-degradative proteinases or proceeding independently of proteolytic remodeling by alternatively using mechanical force to either modulate cell shape or actively distort the fabric of the pericellular ECM (2, 71, 72). As the transcriptional programs that underlie proteolytic versus non-proteolytic cancer cell invasion remain largely undefined, we began our analyses by segregating a single cancer cell line into two discrete populations dominated by either mesenchymal-like collective or amoeboid patterns of invasion. Unexpectedly, the motile fractions of each of these distinct phenotypes expressed a similar cohort of matrix-degrading metalloproteinases. Moreover, both mesenchymal and amoeboid invasion were accompanied by the formation of matrix-cleared tunnels in tandem with the detection of type I collagen degradation products pathognomonic of MMP-mediated collagenolytic activity (35). Consistent with these findings, both mesenchymal and amoeboid -type invasive activity – either in 3D or 2.5D culture – were blocked completely by pan-specific MMP inhibitors. Further, the MMP-dependent, collagen-invasive activity was confirmed in carcinoma cell lines that spontaneously display a mix of both mesenchymal and amoeboid activity (29–31). Finally, despite the fact that cancer cells mobilize a complex mix of MMPs as they infiltrate the surrounding collagen matrix, MMP14/MT1-MMP was identified as the dominant, if not sole, pericellular collagenase driving invasive activity. Of note, while MMP14-targeted cells were rendered immobile in dense type I collagen matrices, the cells do not simply assume a ‘resting’ state. Real-time imaging of knockout spheroids displays brisk intraspheroid trafficking in concert with the upregulation of cell-matrix transcripts normally associated with invasion. As expected, however, proliferative responses are downregulated in tandem with changes in transcripts associated with chromosome segregation, an effect consistent with a requirement for cell ‘pushing’ of the surrounding matrix to engage mitotic machinery (73).

The inability of cancer cells to display collagen-invasive activity in the presence of MMP inhibitors or following MMP14 targeting appears inconsistent with i) the oft-described ability of neoplastic cells to undergo a ‘mesenchymal-amoeboid transition’ following MMP inhibition (1, 15, 16, 20, 40, 70, 74), thereby allowing cells to adopt a non-proteolytic strategy for negotiating matrix barriers, ii) the recent characterization of a mechanical process, termed ‘worrying’, whereby cancer cells physically tear collagen fibrils in the absence of collagenolytic activity (8, 31) or iii) the ability of cells to mobilize a ‘nuclear piston’ to traverse mechanically plastic nanoporous structures (72). First, with regard to an amoeboid-mesenchymal switching mechanism, the original model describing this process was subsequently revised with the discovery that cancer cells are unable to adopt an amoeboid phenotype when encountering rigid matrix barriers with pore cross-sections smaller than ∼7µm^2^ unless proteolytic remodeling was engaged (14). Instead, cancer cells were only able to penetrate size-restrictive ECM barriers via proteinase-independent mechanisms when matrix pore sizes exceeded the limiting dimensions of the rigid cell nucleus (14). In this regard, recent studies have elected to assess proteinase-independent invasion programs utilizing pepsin-extracted type I collagen to provide a matrix with sufficiently large pores (12, 13, 18, 20, 38, 40, 44, 75). Using this non-physiologic material as the starting product, the removal of the collagen N- and C-terminal telopeptides normally found in collagen *in vivo* slows fibrillogenesis, thereby allowing for the generation of matrices with larger pore sizes relative to telopeptide-intact collagen (14, 76). Indeed, using these hydrogels, we find that cancer cell invasive activity is unaffected by either MMP inhibitors or following MMP14 targeting. While these results are consistent with claims that proteolytic requirements for matrix-invasive activity can be bypassed if pore size is sufficiently large, the absence of telopeptides also affects the mechanical properties of the assembled hydrogels (77) Whereas native collagen hydrogels behave as viscoelastic solids, pepsin-extracted collagen hydrogels are devoid of covalent crosslinks and assume plastic characteristics (77, 78). Indeed, when we increased the pore size of hydrogels reconstituted from telopeptide-intact collagen to match that of telopeptide-deleted collagen, proteinase-independent invasion could no longer be maintained. Hence, without increasing mechanical plasticity, even pore sizes far in excess of nuclear dimensions cannot efficiently support proteinase-independent invasion. Unlike manufactured microchannels (47, 79, 80), pores in collagen hydrogels are not aligned in a registered fashion, thereby precluding the spontaneous assembly of long, matrix-free tunnel tracts and necessitating proteolytic remodeling.

Second, a membrane blebbing process has recently been described that allows melanoma cells to invade 3D hydrogels by mechanically fragmenting collagen fibers (8, 31). This result is surprising, because in our work, neither amoeboid fibrosarcoma nor breast carcinoma cells displayed similar activity. It is also worth noting that the bulk of the experiments reported by Driscoll et al. were performed in mechanically plastic, telopeptide-deleted collagen hydrogels (8). Moreover, while a limited subset of experiments in this report (8) and others (20, 81) describe proteinase-independent invasion in telopeptide-intact collagen, we have found marked variability in the levels of collagen covalent cross-linking in commercially obtained preparations (manuscript in preparation). Despite the caveats associated with the use of telopeptide-deleted collagen, we find that MMP14 knockout cancer cells retain full invasive activity in pepsin-extracted collagen hydrogels. Hence, though MMP14 has previously been linked to an array of functions, including the control of cancer cell motile responses, actin-dependent force generation, matrix-adhesive interactions and signal transduction cascades (82–85), 3D invasion programs by MMP14 knockout cells are unaffected in mechanically plastic hydrogels. While recent studies have described the ability of normal as well as neoplastic cells to create migration paths in viscoelastic and/or plastic matrices via mechanical processes alone (8, 72, 77), neither native collagen hydrogels nor tumor tissue explants display the physical properties necessary to support proteinase-independent invasion. Importantly, our results should not be construed to suggest that specific tissues (e.g., the proteoglycan/glycosaminoglycan-rich brain matrix (86)) may not be permissive for proteinase-independent invasion, but rather that type I collagen-rich tissues require the mobilization of MMP14, a finding consistent in our recent studies of breast cancer invasion *in vivo* where metastatic activity was obviated when carcinoma cell MT1-MMP was deleted specifically (87). We also note that a number of cell intrinsic and extrinsic microenvironmental factors, ranging from changes in nuclear rigidity, chemoattractant gradients and media viscosity to ECM composition and collagen fibril alignment have been reported to modulate cancer cell invasion programs (47–58). In our studies, however, none of these elements restored invasive activity to MMP14 knockout cells in intact collagen hydrogels. Perhaps most surprising in these efforts is the fact that CAFs, though capable of promoting the invasive activity of wild-type cancer cells and degrading pericellular type I collagen, did not confer invasive activity to MMP14 knockout cancer cells. However, previous reports i) did not examine the ability of wild-type CAFs to direct the invasive activity of MMP14 knockout cancer cells (50, 55–57), ii) alternatively generated hybrid spheroids or cell mixtures that contain both cancer cells and CAFs (55, 58), a construct that may not accurately recapitulate cancer cell-stromal cell interactions *in vivo* or iii) used collagen-Matrigel hydrogels (55, 57, 58) where Matrigel interferes with type I collagen fibrillogenesis while displaying highly plastic mechanical properties in its own right (88, 89). Further, wild-type cancer cell spheroids cannot rescue the invasive activity of juxtaposed MMP14 knockout cells, ruling out the possibility of exosome-dependent MMP14 transfer or the release of MMP14-derived pro-motility factors (85, 90). Nevertheless, wild-type cancer cells can, as previously reported (91), rescue the invasive activity of MMP14 null cancer cells when co-mixed as hybrid spheroids, likely by creating collagen-free tunnels that motile-active knockout cells can then capitalize.

As matrix concentration increases, matrix pore size and rigidity as well as ligand density change in a coupled fashion. A number of elegant bioengineering tools using synthetic hydrogels have been developed in an effort to deconvolute the relative effects of each of these physical variables on cell behavior (92–96). *In vivo*, however, these properties do not change in singular fashion. As type I collagen levels increase or decrease in pathophysiologic settings, attendant changes in matrix density, pore size and rigidity occur in a mechanically coupled manner, rendering an experimental approach where the collagen concentration is varied *in vitro* most relevant to the *in vivo* setting (93). Indeed, though a body of evidence supports the proposition that increased matrix rigidity promotes aggressive characteristics in cancer cells (60–62, 97), we and others find that cancer cell invasive potential decreases markedly as collagen concentration increases, a result consistent with reports that type I collagen acts as a barrier to the spread of neoplastic cells *in vivo* (98–101). By contrast, while decreasing collagen concentration augmented invasive activity in our studies, it more importantly provided our first example of MMP14-independent invasion through a crosslinked matrix barrier. Though the large pore sizes found in type I collagen hydrogels prepared at a final concentration of 1 mg/ml might have been predicted to obviate a requirement for MMP14 expression or activity (43, 102), invading cells were consistently found in association with proteolyzed collagen fragments. In this scenario, migratory capacity was maintained in the absence of MMP14-mediated collagenolysis, but only at the expense of nuclear integrity where we posit that that the distortions of cell shape required to circumnavigate the rigid collagen fibrils damaged the rigid nuclear envelope. Whether nuclear rupture ultimately undermines cancer cell function or serves to promote genetic changes capable of engendering metastatic behavior (79, 80, 103, 104), will be the subject of future study. Of note, reports have described loss of lamin B1 and accrual of DNA damage in MMP14-targeted cells in 2D (105, 106), however we do not detect similar changes in our MMP14-targeted cells under these conditions (Supplementary Figure 16a and b).

Finally, while matrix constructs have provided invaluable information on cancer cell invasion programs, *in vitro* models cannot recapitulate the complex stromal cell-ECM environment encountered *in vivo* (10). Using live human tissue explants recovered from breast cancer patients, we confirmed a critical role for MMP14 in conferring breast carcinoma cells with the ability to infiltrate the type I collagen-rich interstitial matrix. We do note that occasional areas of invasion were observed with MMP14 knockout carcinoma cells or even in the presence of pan-specific MMP inhibitors. However, collagen concentrations may vary in association with neoplastic sites and the deposition of other ECM components, ranging from fibronectin and laminin to fibrin, that are sensitive to a wide range of proteolytic enzymes must be considered (107, 108). Likewise, the ability of cancer cells to locate matrix-poor zones that allow proteinase-independent infiltration over limited distances cannot be ruled out and may provide an explanation for amoeboid-like movement observed in short term, real-time imaging of tumor sites *in vivo* (15, 109, 110). However, caution should be exercised in terms of ascribing proteinase-free cancer cell movement to earlier descriptions of large, collagen-free pores observed *in vivo* as assessed by second harmonic generation imaging (111, 112). While this technique can resolve larger diameter type I collagen fibers, small diameter fibrils remain invisible as do most ECM macromolecules save for type I-III collagens (42, 113). Indeed, using nanoparticle diffusion to assess *in vivo* matrix properties of malignant tissues, pore sizes have been reported to be even smaller than those typically observed in collagen hydrogels generated *in vitro* (93). In this regard, recent studies demonstrate that fast moving breast cancer cells observed *in vivo* are devoid of metastatic potential relative to slower moving matrix-degradative carcinoma cells (114). While these data, taken together with our own, support a key, if not dominant, role for MMP14 in regulating tissue-invasive behavior in cancer, they fall short of implicating an MMP14-type I collagen interactome that operates in the setting of human disease. As such, we leveraged a spatial transcriptomic dataset to probe human cancer lesions for *MMP14* expression, specifically within invasive regions. These data revealed a strong association between *MMP14* expression and invading tumor cell populations. Of note, MMP14 was not limited only to tumor cells, and could also be detected, though with less consistency, in other annotated areas containing stromal and immune populations. These data were not unexpected, as multiple cell populations, including fibroblasts (115), endothelial cells (116) and macrophages (117), are capable of expressing MMP14. As these data were limited to MMP14 at the transcript level, we finally performed immunohistological analyses on human breast cancer tissues, finding that invasive lesions are decorated with ¾ collagen degradation products that are specific for MMP-dependent collagenolytic attack. Interestingly, our clinical samples included cases of invasive lobular carcinoma whose single-file pattern of tissue infiltration is likened to an amoeboid, rather than collective, pattern of invasion (69). While clinical examples of invasive lesions in association with matrix degradative signatures are rarely described, Karin and colleagues recently reported that type I collagen degradation products are found in association with human pancreatic cancer (118), but it should be noted that the staining pattern detected in their work is more focal and does not display the striated pattern we observe in association with invading strands of breast cancer cells. Further, in the case of breast cancer, type I collagen degradation products are found in association with MMP14 expression at the protein level, but only in areas devoid of an intervening basement membrane. Similar to our findings here, we have previously described the constitutive expression of epithelial Mmp14 in the normal mouse mammary gland (87), but the proteolytic activity of the proteinase (i.e., its expression in latent vs active form and the level of endogenous inhibitors) has not been previously determined. However, as active MMP14 can degrade the basement membrane (117), it seems most likely that the proteinase is either confined to intracellular compartments, exocytosed in latent form or complexed with endogenous inhibitors (119). In any case, though an association between MMP14 expression and type I collagen degradation does not prove causation, our findings *in vivo* coupled with our *in vitro* analyses support the contention that MMP14 is essential for cancer cell invasion programs operative in type I collagen-rich stromal tissues and that while cells do exhibit mesenchymal-amoeboid transition, amoeboid-type invasion remains reliant on MMP14 activity. The failure of MMP inhibitors to exert palliative effects in clinical trials is often used as the strongest evidence for proteinase-independent invasion. However, the specificity and efficacy of targeting matrix-degradative enzymes *in vivo* has never been defined or proven (120, 121). While inhibiting MMP14 activity specifically in select cell populations may not be feasible, identifying the transcriptional programs that drive its expression or the processes that control its exocytosis to the cell surface are deserving of careful consideration.

## Materials and Methods

### Cell culture

HT1080, MDA-MB-231, SUM159 and HEK293T cells were all acquired through ATCC and maintained in complete media, high glucose DMEM (Gibco, 11995-0665) with 10% fetal bovine serum (Corning, 35-015-CV) and 1x Penicillin/Streptomycin (Gibco, 15140-122), under standard culture conditions (5% CO_2_/air, 37°C). Human breast cancer-associated fibroblasts, isolated as described (59), were maintained in modified IMEM (Gibco, A1048901) with 20% FBS, on gelatin-coated (Cell Biologics, 6950) plates. Cells were routinely confirmed to be negative for mycoplasma using the LookOut Mycoplasma PCR Detection Kit (Sigma, MP0035). HT1080 and MDA-MB-231 cells transiently expressing Life-Act-GFP (Addgene, #58470) were prepared using the Lipofectamine 2000 Transfection Reagent (Invitrogen, 11668027). Stable GFP-NLS expressing lines were prepared through lentiviral transduction. Briefly, HEK293T cells were transfecting using Lipofectamine 2000 along with GFP-NLS plasmid (Addgene, 132772) and packaging plasmids, psPAX2 (Addgene, 12260) and pMD2.G (Addgene, 12259) for 6 hours. Following transfection, media was replaced, and cells were cultured for 24-48 hours. Virus containing media was isolated and filtered through a 40µm filter before being applied to cells in the presence of polybrene (Sigma TR-1003) for transduction. Following several passages, cells were FACS sorted for green, fluorescent signal.

### 3D spheroid invasion assay

Spheroids were prepared using the hanging droplet method (122), whereby single-cell suspensions of tumor cells were cultured in DMEM with 0.2% methylcellulose (Sigma, M7027) to support aggregation and plated on the lid of culture dishes humidified with PBS in the main compartment of the dish. Specifically for MDA-MB-231 cells, to further support spheroid formation, 18µg/ml atelocollagen (PureCol) was included in the media (123). HT1080 and SUM159 spheroids were allowed to form for 2-3 days, while MDA-MB-231 spheroids were generated during a 5–6-day culture period. Mixed control and MMP14-KO spheroids were prepared in the same way with a ratio of 1:10 control to MMP14-null tumor cells.

Acid solubilized rat tail collagen was prepared as previously described (124). Briefly, rat tail tendons are stripped from the tail and transferred to 70% ethanol. Tendons are then pulled apart into smaller fibers and allowed to dry before undergoing collagen extraction in 0.2% acetic acid. After extraction, the solution is centrifuged at 30,000g for 30 min. Supernatant is collected and lyophilized. Lyophilized collagen is then weighed and re-dissolved in 0.2% acetic acid. Collagen hydrogels were prepared on ice by mixing either in-house prepared acid-solubilized rat tail collagen or pepsin-extracted bovine collagen in 1M HCl (PureCol, Advanced Biomatrix, 5005) with 10x MEM (Gibco, 11430030), 1M HEPES (Gibco, 15630080) and 0.34N NaOH to reach the desired final concentration of collagen, with 1x MEM, 0.025M HEPES and final pH of 7.4. Upon neutralization of the collagen solution, on ice, formed spheroids were washed off of the culture dish lids and resuspended in collagen solution which was then plated and allowed to polymerize at either 37°C or 8°C. After hydrogel polymerization, complete media was added atop the gel, and spheroids allowed to invade for ∼72 hours.

For all treatments, additive/inhibitor concentrations were initially prepared accounting for both the gel and media volumes in the assay and media was replaced every 2 days. MMP inhibition was performed using BB-94 (Batimastat, 5µM, Tocris, 29-611), GM6001 (Illomastat, 10µM, Selleckchem, S7157), TIMP-1 (12.5µM, Gibco, 410-01) and TIMP-2 (5µM, Gibco, 410-02). Media additives included the anti-ß1 integrin antibody, 4B4 (2µg/ml, Beckman Coulter, 6603113), SDF1 (100ng/ml, Gibco, 300-28A), EGF (10ng/ml, Gibco, AF-100-15) and 15cp methylcellulose at 1% for a final viscosity of ∼8cp. Low nutrient conditions included low glutamine (0.2mM), low glucose (0.5mM) and hypoxia (0.2% O_2_, 5% CO_2_ with N_2_ balance). Hydrogel additives were added to neutralized collagen solutions prior to polymerization and included type VI collagen (50µg/ml, Abcam, ab7538) and fibronectin (25µg/ml, Sigma, F1141). For tension-based alignment of collagen fibers, collagen hydrogels were allowed to polymerize with two pins from an ‘alignment apparatus’ (Figure 6a) inserted in the collagen, roughly 6mm apart from one another. Following polymerization, using a screw based expanding mechanism, the pins were expanded to separate an additional 2mm apart, generating a region of tension-aligned collagen between the two pins. Spheroid and cancer-associated fibroblast co-culture experiments were performed by adding CAFs at 500,000 cells/ml to the collagen suspension used to resuspend tumor spheroids.

### 2.5D collagen invasion

Neutralized type I collagen solutions were prepared as described for 3D spheroid invasion assays and plated on Transwell filters with 3µm pore size (Steriltech, 9323002). Following collagen polymerization, cell suspensions of 150µl at 2.5X10^5^ cells/ml in complete media were added atop the collagen hydrogels in the upper chamber and complete media alone was additionally added to the bottom chamber. Cells were allowed to invade into the collagen hydrogel for 6 days under standard culture conditions.

### Transwell and 2D migration assays

For Transwell migration, single-cell suspensions were added atop 24-well Transwell inserts with 3µm or 8µm pore sizes (Steriltech, 9323002 & 9328002) with complete media in the top and bottom chamber. After allowing cells to adhere overnight, the top media was replaced with serum-free high glucose DMEM and cells were allowed to migrate for 24 hours using an 8µm pore or 48 hours for 3µm pore filters. Transwells were fixed in 70% ethanol, the upper surface of the filter was swabbed to remove cells that had not transmigrated. Cells that had crossed the filter surface were stained with Crystal Violet, imaged and invading cells counted.

For 2D motility assays, cells were plated at high confluence in 48-well plates in complete media and allowed to adhere overnight. The following day, a single scratch was made through the center of each well. Following scratch generation, cells were washed with serum-free DMEM and imaged. Serum-free DMEM was added to the samples and cells were allowed to migrate overnight before being re-imaged.

### Cell proliferation assay

Cells were seeded on day 0 in their respective maintenance culture media in 24-well plates at a density of 10,000 cells per well. After four days of culture, the cells were trypsinized and counted. The cell numbers of the knockout cell lines were normalized to those of their respective control cell lines. Media were replenished every 24 hours.

### Fluorescent collagen labeling and assessment of alignment

Un-labelled collagen hydrogels were prepared in a 10cm plate. Following polymerization of the collagen, a fluorescently tagged NHS-Ester dye (Sigma, 41698) was used to label the collagen overnight at 4°C. The labelled collagen was then washed with DPBS before being re-solubilized in 500mM acetic acid overnight with agitation. After the labelled collagen hydrogel was solubilized, the collagen solution was dialyzed against 0.2% acetic acid. Final collagen concentration was calculated using the Sircol 2.0 soluble collagen assay (BioColor, SIRC2) following the manufacturer’s instructions. Final working solutions of fluorescently labelled collagen were prepared by mixing labelled and unlabeled collagen to reach 8% labelled collagen. Collagen alignment was assessed using the OrientationJ plug-in in ImageJ (v2.16.0/1.54n).

### PEG hydrogel formulation and invasion

Poly(ethylene glycol)-norbornene (PEGNB) hydrogels were synthesized in a one-pot synthesis via thiol-ene photopolymerization. 4-Arm PEGNB (20kDa; PSB-4112, Creative PEGWorks, Chapel Hill, NC) and lithium phenyl-2,4,6-trimethylbenzoylphosphinate (“LAP”; 900889, Sigma-Aldrich, St. Louis, MO) were sourced from commercial suppliers that provide the percent substitution of norbornene by NMR and purity by HPLC, respectively. The thiol-containing cell-adhesive peptide Ac-CGRGDS-NH2 (“RGD”; AAPPTEC, Louisville, KY) and dithiol-containing matrix-metalloproteinase-(MMP-) sensitive crosslinking peptides Ac-GCRDVPMS↓MRGGDRCG-NH2 (“VPMS”, cleavage site indicated by ↓; AAPPTEC, Louisville, KY), which feature N-terminal acetylation and C-terminal amidation, were dissolved in 25 mM acetic acid (A38-212, Fisher Scientific, Hampton, NH), filtered through 0.22 µm filters (SLGV013SL, Sigma-Aldrich, St. Louis, MO), lyophilized for 2 days, and stored in a desiccator at -20 °C. The purity of peptide aliquots was determined using Ellman’s reagent (22582, Thermo Fisher, Waltham, MA) per the manufacturer’s protocol. PEGNB, LAP, and MMP-insensitive crosslinker PEG-dithiol (PEGDT, 3400 Da; Laysan Bio, Inc., Arab, AL) were resuspended in phosphate-buffered saline (PBS, pH 7.4, 1X; 10010-023, Gibco, Waltham, MA) and sterile filtered with 0.22 µm filters to prepare fresh stocks for each experiment. Peptides were resuspended in PBS to reach desired stock concentrations. To create a stock solution, 3% wt. PEGNB (w/v), 1 mM LAP, 1 mM RGD, 90% VPMS or PEGDT (added to theoretically achieve 0.9 thiols per norbornene after accounting for by RGD incorporation), and PBS were combined together to resuspend spheroids. The stock solution was gently vortexed, and 50 µL of solution was transferred into 1 mL syringes (14-823-434, Fisher Scientific, Hampton, NH) with the tips cut off. Syringes were placed ∼1 inch under a 6-Watt LED 365 nm Gooseneck Illuminator (LED-6W-UV365, AmScope, Feasterville, PA) for 90 seconds at maximum UV light, corresponding to approximately 50 mW/cm2. Hydrogels were subsequently placed into 2 mL of media and spheroids were allowed to invade for 4 days prior to fixation.

### Rheology

Hydrogel shear storage (G’) moduli were measured one day following hydrogel polymerization. Hydrogels were gently removed from their wells and centered between a Peltier plate and an 8-mm measurement head on an AR-G2 rheometer (TA Instruments, New Castle, DE). To reduce slippage during measurement, the plate and measurement head were covered with P800 sandpaper (599-18-CC, 3M, Saint Paul, MN). The measurement head was lowered until it touched the hydrogel, then it was lowered an additional 300 µm. After a 30 second pause for temperature equilibration, G’ and G” were averaged over a 1 min time sweep, measured at 37 °C, 5% strain amplitude, and 1 rad/s frequency.

### Single-cell RNA-sequencing

Spheroids were prepared at 2,000 cells per droplet, and ∼50 spheroids per sample were embed in 500µl collagen, allowing for 72 hours of 3D spheroid invasion. At this time, media were aspirated, and collagen hydrogels were gently removed from their well and transferred to a 1.7ml tube. Using small surgical scissors, gels were cut into pieces before adding sterilized digestion buffer (2mg/ml Type 7 Collagenase (Worthington) in HBSS) at a 2:1 buffer to gel ratio. Samples were incubated at 37°C for 10 minutes before lightly vortexing and pipetting the samples. After an additional 5-10 minutes, when collagen was no longer visible, cells were pelleted in a microcentrifuge at 300g. Digestion buffer was aspirated, and the isolated cells were resuspended in 100ul trypsin (0.25%) and incubated at 37°C for an additional three minutes before 1ml of complete media was added to inhibit proteolysis. Samples were filtered into a 5ml falcon tube with a cell strainer cap before being pelleted by centrifugation (300g). Media was aspirated and cells were resuspended in 50µl FACS buffer (PBS + 0.5% BSA) for sequencing.

Prepared single-cell suspensions were used for library preparation with 10X Genomics Chromium Single-Cell 3’ v3 Chemistry following manufacturer’s protocols. Libraries were subjected to 151bp paired-end sequencing using Illumina NovaSeq to a depth of ∼60,000-150,000 reads/cell and BCL Convert Conversion Software (v4.0, Illumina) was used to generate de-multiplexed FASTQ files. Sequencing data were processed using the 10X Genomics CellRanger pipeline (7.1.0) and align to the GRCh38-2020-A human reference genome to generate gene expression counts. Following alignment and initial quality control, 3478 and 1567 cells from control and 4B4-treated HT1080 spheroids, respectively, were included in the analysis using the Seurat package (5.0.1) in R (4.3.2). Following additional filtering for cells with 12000 > nFeature_RNA > 3000 and percent mitochondrial RNA < 17.5 percent, 3327 and 1273 cells were included in the final analysis using Seurat. Gene sets used for transcriptomic characterization are included in Supplementary Table 1.

### RNA sequencing

MDA-MB-231 spheroids were generated (2000cell/spheroid, 100,000 total cells/sample) and RNA was prepared either following 5 days of spheroid formation in hanging droplets, 24 hours post-embedding in collagen hydrogels or 48 hours post-embedding in collagen hydrogels. RNA samples were extracted using RNeasy Mini Kit (Qiagen, 74104), following collagen gel digestion as described for single-cell isolation, without the trypsinization or filtering steps. RNA-seq libraries were prepared using standard Poly-A Library Prep (Illumina) and subjected to 151bp paired-end sequencing according to manufacturer’s protocols (Illumina NovaSeqXPlus). BCL Convert Conversion Software (v4.0, Illumina) was used to generate de-multiplexed FASTQ files. Reads were trimmed using Cutadapt (v2.3) (125). FastQC (v0.11.8) (126) was used to ensure the quality of data. Fastq Screen (v0.13.0) was used to screen for various types of contamination (127). Reads were mapped to the reference genome GRCh38 (ENSEMBL), using STAR v2.7.8a (128) and assigned count estimates to genes with RSEM (v1.3.3) (129). Alignment options followed ENCODE standards for RNA-seq. Multiqc (v1.7) compiled the results from several of these tools and provided a detailed and comprehensive quality control report (130). Samples had an average of ∼35 million reads each and were analyzed in R (4.3.2) using DESeq2 (1.42.0). DEGs were defined as genes with adjusted p-value < 0.05 and fold change greater than |1.5|. Full DEG lists for MMP14-KO versus control samples at each timepoint are available in Supplementary Table 2. Analysis of large-scale bulk RNA sequencing datasets were performed using TNMplot.com (65).

### Generation of MMP14-mCherry knock-in

Left and right homology arms were inserted into the pDonor-D08 plasmid (GeneCopoeia, GEN-pDonor-D08) with the right arm containing the mCherry tag inserted between amino acids 515 and 516 on the MMP14 gene. Guide RNAs (hMT1-sg1-F: CAC CGG ATG AAT GGG CAC CAC GTG and hMT1-sg1-R:AAA CCA CGT GGT GCC CAT TCA TCC; hMT1-sg2-F: CAC CGA CGG AGG TGA TCA TCA TTG and hMT1-sg2-R:AAA CCA ATG ATG ATC ACC TCC GTC were incorporated into the pX330-U6-Chimeric_BB_CBh-hSpCas9 plasmid (Addgene, 42230). Cells were co-transfected with the donor and guide RNA plasmids using Lipofectamine 2000 and selected with G418 for ∼10 days. Cells were then transfected with a Puro-Cre plasmid (Addgene, 17408) to flox out the GFP and Neomycin reporter cassettes. Cells were then FACS sorted to remove residual GFP expressing cells with un-floxed reporter sequences.

### CRISPR/Cas9 gene targeting

Guide RNAs were designed using CHOP-CHOP (131) and ligated into the LentiCrispr-V2-mCherry backbone (Addgene, 99154). HEK293T cells were transfecting using Lipofectamine 2000 along with LentiCrispr plasmids containing the guide RNA and packaging plasmids, psPAX2 and pMD2.G for 6 hours. Following transfection media was replaced and cultured for 24-48 hours. Virus containing media was isolated and filtered through a 40µm filter before being directly applied to cells in the presence of polybrene (Sigma, TR-1003) for transduction. Following several passages cells were FACS sorted for mCherry, fluorescent signal. FACS-sorted bulk knock-out populations were used for experiments to avoid confounding clonal differences. MMP14 was targeted using two guide RNAs with forward sequences of hMMP14-sg1-F: CAT CCG GCC GGC CTC CCG AT and hMMP14-sg2-F: GAT GCC CCG GCG GTC ATC AT. LMNA was targeted using hLMNA-sg1-F: AGT TGA TGA GAG CCG TAC GC.

### Human breast tissue invasion and histology

De-identified human breast tissue samples were harvested immediately following surgery, suspended in RPMI 1640 (Gibco, 11875135) with penicillin and streptomycin, and shipped on ice by The Cooperative Human Tissue Network (CHTN) to the laboratory. A summary of tissue pathology can be found in Supplementary Table 3. Upon receipt, portions of the samples were immediately fixed in 10% neutral buffered formalin overnight before being processed for paraffin embedding. The remaining live tissue from normal breast samples was sliced and plated atop a small 25µl droplet of collagen solution in a Transwell, used to anchor the tissue sample to the Transwell filter. Care was taken to ensure the upper surface of the tissue slice did not come into contact with the collagen solution so cells seeded atop the explant would directly interface with the human tissue. Cells were then seeded in the upper chamber of the well atop the tissue slice at 10^5^ cells/ml. Complete media containing 5ng/ml EGF (Gibco, AF-100-15) was added to the bottom chamber. Cells were allowed to invade for 6 days. Samples were then fixed in 4% PFA. Fixed samples were embedded in paraffin on edge and sectioned. H&E and immunofluorescence staining was performed on paraffin sections.

### Spatial Transcriptomics

We obtained the human breast cancer Spatial Transcriptomics dataset (66) from https://github.com/almaan/her2st and analyzed all 8 samples (A1, B1, C1, D1, E1, F1, G2, H1) with pathologist’s annotations. The R package STutility (v. 1.1.1) was used to load the dataset into R (v. 4.4.0). Following established quality control settings (66), genes expressed in fewer than 3 spots and spots with expression of fewer than 100 genes were filtered out (Supplementary Table 4). UMI count matrices were normalized using the *NormalizeData* function from Seurat (v. 5.1.0). Specifically, the UMI counts for each spot were divided by the total counts for that spot, multiplied by a scaling factor of 10,000 and the results log-transformed using a pseudo-count of 1. The metagene consisting of MMP14 and KRT7 was defined as the sum of normalized expressions following a previously described method (132). The Z-scores were obtained by standardizing the normalized expressions to have 0 mean and unit variance.

### Western blotting

Cell lysates were prepared using Pierce RIPA buffer (ThermoFisher, 89900) containing 1x Pierce Protease Inhibitor (ThermoFisher, A32953). DNA was sheared via passage through a 1 ¼ inch 26-gauge needle ∼6 times. Protein concentration was determined using the Pierce BCA Protein Assay Kit (ThermoFisher, 23225, 23227, A65453) following manufacturer’s instructions. Extracts were analyzed using standard techniques with horseradish peroxidase-conjugated secondary antibodies (Cell Signaling) and detected by the Pierce SuperSignal West Pico system (ThermoFisher, 34577). Membranes were cut as indicated in final unprocessed images in order to use different primary antibodies on the same blot for loading controls. Original blot images can be found in Source Data. Additional antibody information is available in Supplementary Table 5.

### Immunofluorescence/Immunocytochemistry and confocal reflectance imaging

*In vitro* samples were fixed in 4% paraformaldehyde (Electron Microscopy Services, 15714-S) and washed with PBS and blocked for 1 hour prior to primary antibody staining overnight at 4°C for 2D samples and 48 hours at 4°C for 3D samples. Samples were washed with PBS before the addition of secondary antibodies (including phalloidin and DAPI) and stained 1 hour at room temperature for 2D samples and overnight at 4°C for 3D samples. After extensive washing with PBS samples were images directly in the chambered coverslips used for the experiments. CellMask-647 (Invitrogen, C56129) and Hoechst 33342 (Life Technologies, H3570) were used following manufacturer instructions.

For staining of human tissue, paraffin sections were rehydrated in a xylene/ethanol series and microwave-mediated antigen retrieval performed in pH 6.0 1M citrate buffer. Sections were blocked for 1 hour in donkey serum prior to the addition of primary antibodies directed against type I collagen, type IV collagen, Pan-cytokeratin, ¾ collagen and MMP14. The specificity of the MMP14 antibody was confirmed by staining paraffin-embedded wild-type or MMP14-knockout MDA-MB-231 cells (Supplementary Figure 19). Samples were incubated overnight at 4°C and washed with PBS before the addition of secondary antibodies for 1 hour at room temperature. Slides were washed again with PBS and stained with DAPI for 15 minutes. Following a final wash, coverslips were mounted using Prolong Gold. Image acquisition was performed using the Nikon Ax Confocal Microscope System with NIS Elements Software (v5.42.06).

### Live-Cell imaging

Image acquisition was performed using a spinning disc confocal CSU-WI (Yokogawa) on a Nikon Eclipse TI inverted microscope and Micro-Manager software (Open Imaging). Live imaging was performed on unfixed cells incubated under standard conditions in a stage-top live cell apparatus (Livecell Pathology Devices). Cell tracking and quantification of invasion speed and change of direction were performed using the TrackMate plug-in in ImageJ. Images were recorded using either brightfield with a 10x objective lens or confocal fluorescence using a 60x oil-immersion lens. For fluorescent images cells were labeled with CellMask-647 (Invitrogen, C56129) and Hoechst 33342 (Life Technologies, H3570).

### Quantification and statistics

All data are shown as mean values with error bars representing the standard error (SEM). Adobe Photoshop, ImageJ (NIH) and GraphPad Prism 7 software were used to process and analyze images and data. For each comparison between two groups, statistical analysis was performed, and p values were calculated with an unpaired two-tailed Student’s t test with Welch’s correction using GraphPad Prism 7 software. For 3D invasion quantification, Invasion Score was calculated as area of all nuclei (invasive and non-invasive) minus the area of non-invasive nuclei (confined to the original spheroid boundaries). For experiments where cellular morphology was comparable between samples, nuclear area could be substituted with total cellular area. Single-cell RNA sequencing comparisons were made using the Wilcoxon Rank Sum Test. For all tests, p<0.05 was considered statistically significant.

## Acknowledgments

Work performed in this study was supported by grants from the National Institutes of Health, R01-CA-071699 (S.J. Weiss), R01-HL-085339 (A.J. Putnam), F31-CA-275102 (A.W. Olson), NCI Training Grant T32-CA-009676 (A.W. Olson), NHLBI T32-HL-125242 (A.J. McCoy), the Breast Cancer Research Foundation (S.J. Weiss), the Margolies Family Discovery Fund for Cancer Research (S.J. Weiss), and the Rogel Cancer Graduate Student Scholarship of the University of Michigan Rogel Cancer Center (A.W. Olson). Library prep and next-generation sequencing was carried out in the Advanced Genomics Core at the University of Michigan. Research reported in this publication was supported by the University of Michigan Advanced Genomics Core, the UM Single-Cell Spatial Analysis Program and the National Cancer Institutes of Health under Award Number P30CA046592 by the use of the following Cancer Center Shared Resource: Single-Cell and Spatial Analysis Shared Resource.

