## Supplementary Figures 1-19 for "Amoeboid-Mesenchymal Transition and the Proteolytic Control of Cancer Invasion Plasticity"

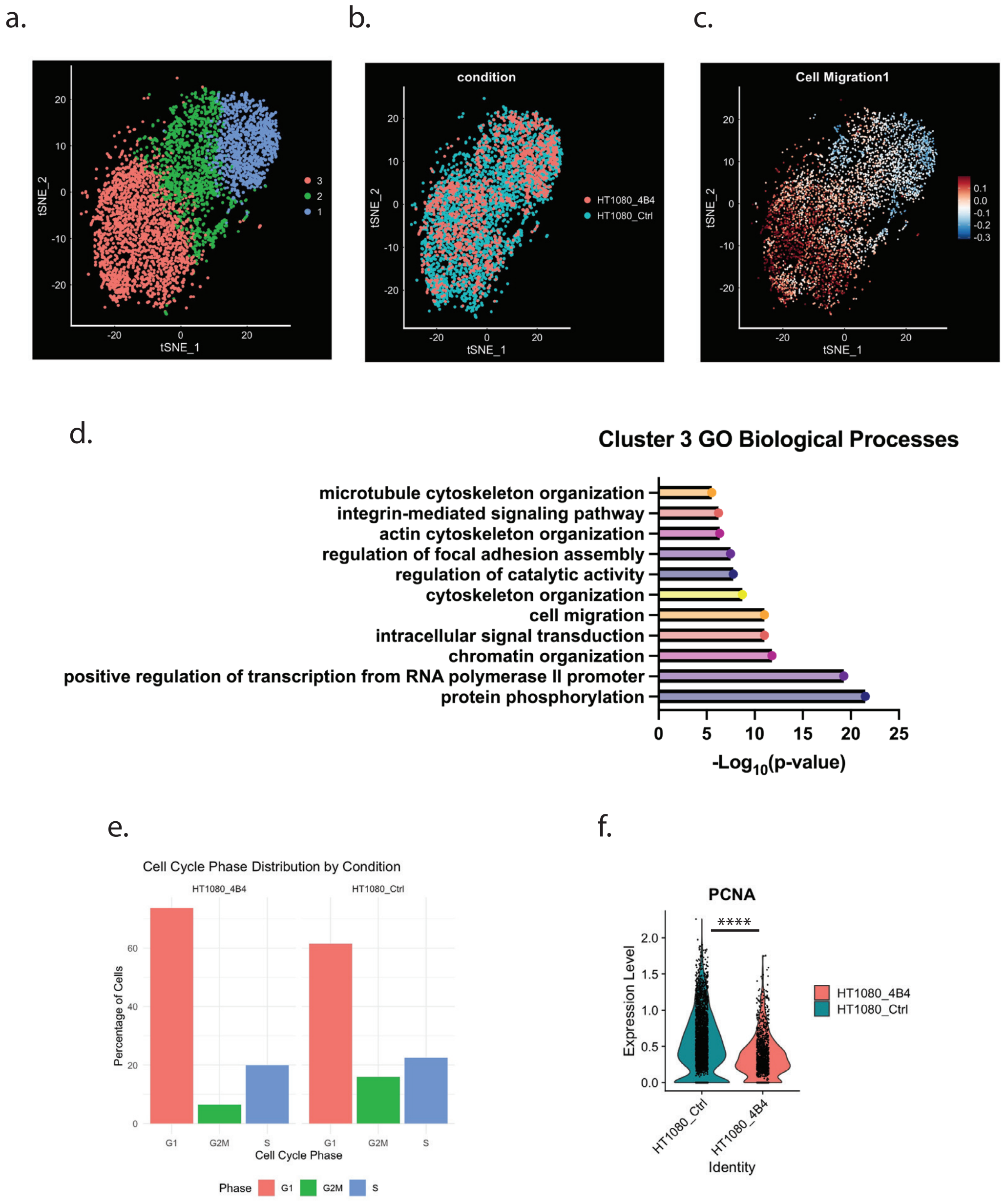


**Supplementary Figure 1: Merged scRNAseq Characterization.** a-c) tSNE plots depicting the clustering of combined samples and their expression of migration-associated genes. d) Gene Ontology biological processes up in control versus 4B4-treated invading cells. Quantification of cell cycle phases detected by transcriptional profiles. f) Violin plot of PCNA expression between control and 4B4-treated samples. P-value < 0.05 (*), <0.01 (**), <0.001 (***), < 0.0001 (****).

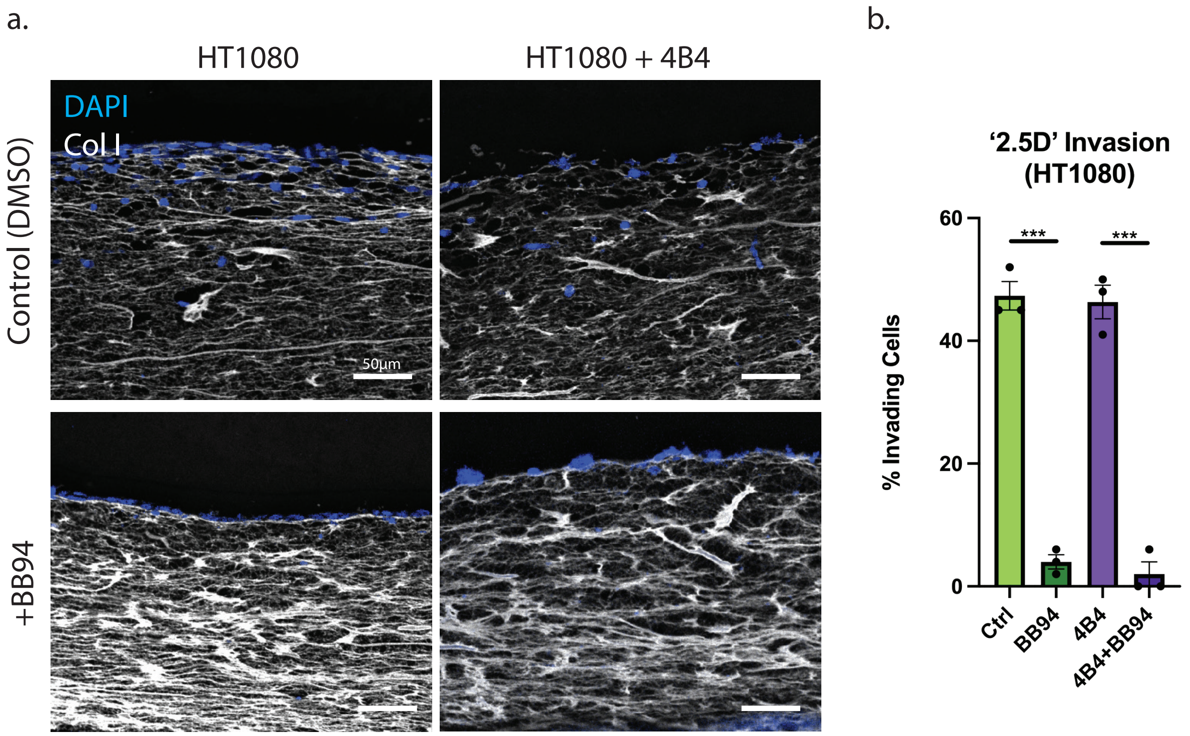


**Supplementary Figure 2: 2.5D Invasion of Collagen Hydrogels.** a) Representative images of '2.5D' invasion of HT1080 cells into underlying 3D collagen hydrogels. b) Quantification of the percentage of cells observed invading into the underlying collagen hydrogel. P-value < 0.05 (*), <0.01 (**), <0.001 (***), < 0.0001 (****).

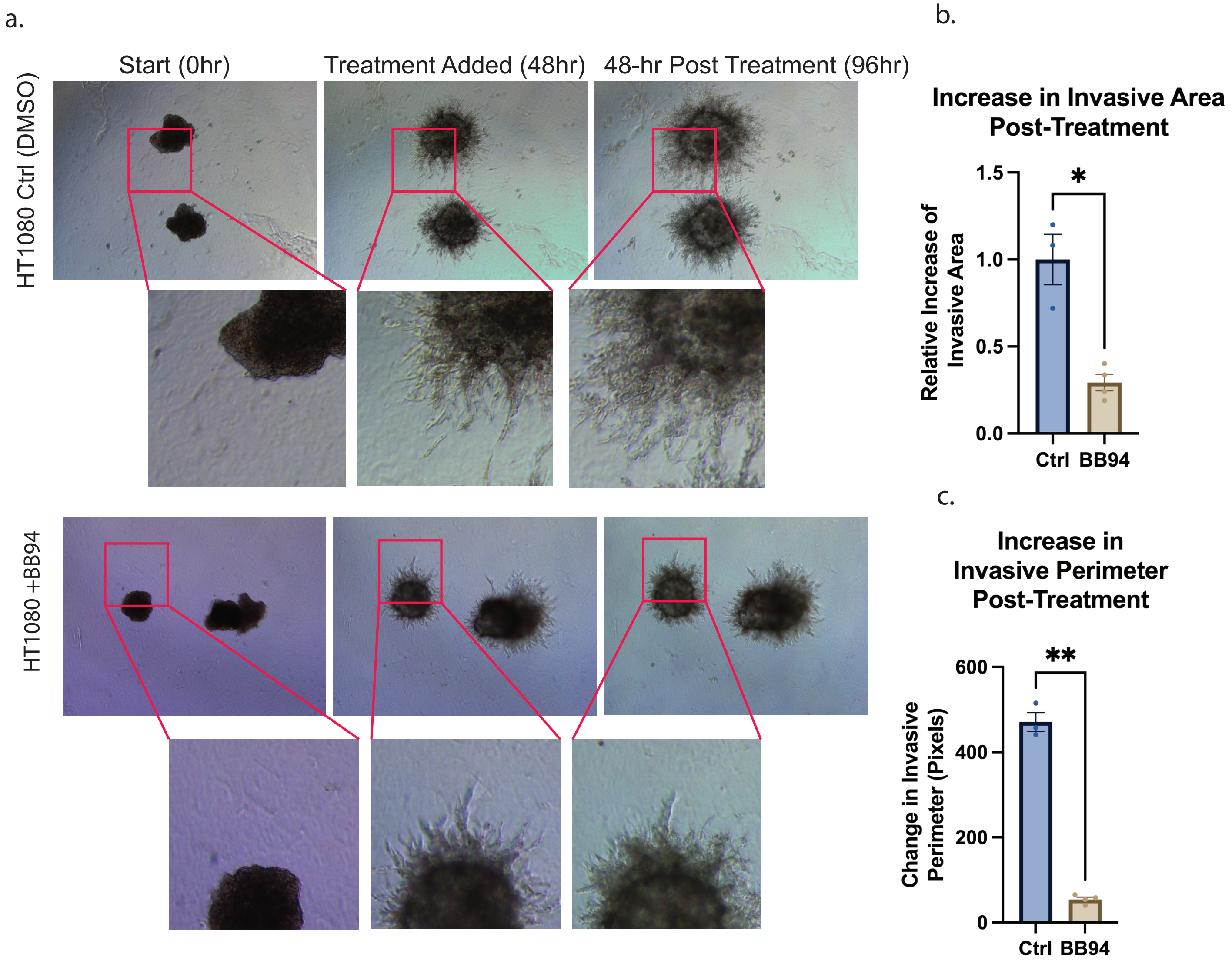


**Supplementary Figure 3: Delayed Addition of MMP Inhibitors Blocks Tumor Cell Invasion.** a) Representative phase-contrast images of spheroid invasion in 3D collagen hydrogels at the indicated time points. b,c) Quantification of the change in invasive area and invasive perimeter in the 48 hours following addition of the treatment. P-value < 0.05 (*), <0.01 (**), <0.001 (***), < 0.0001 (****).

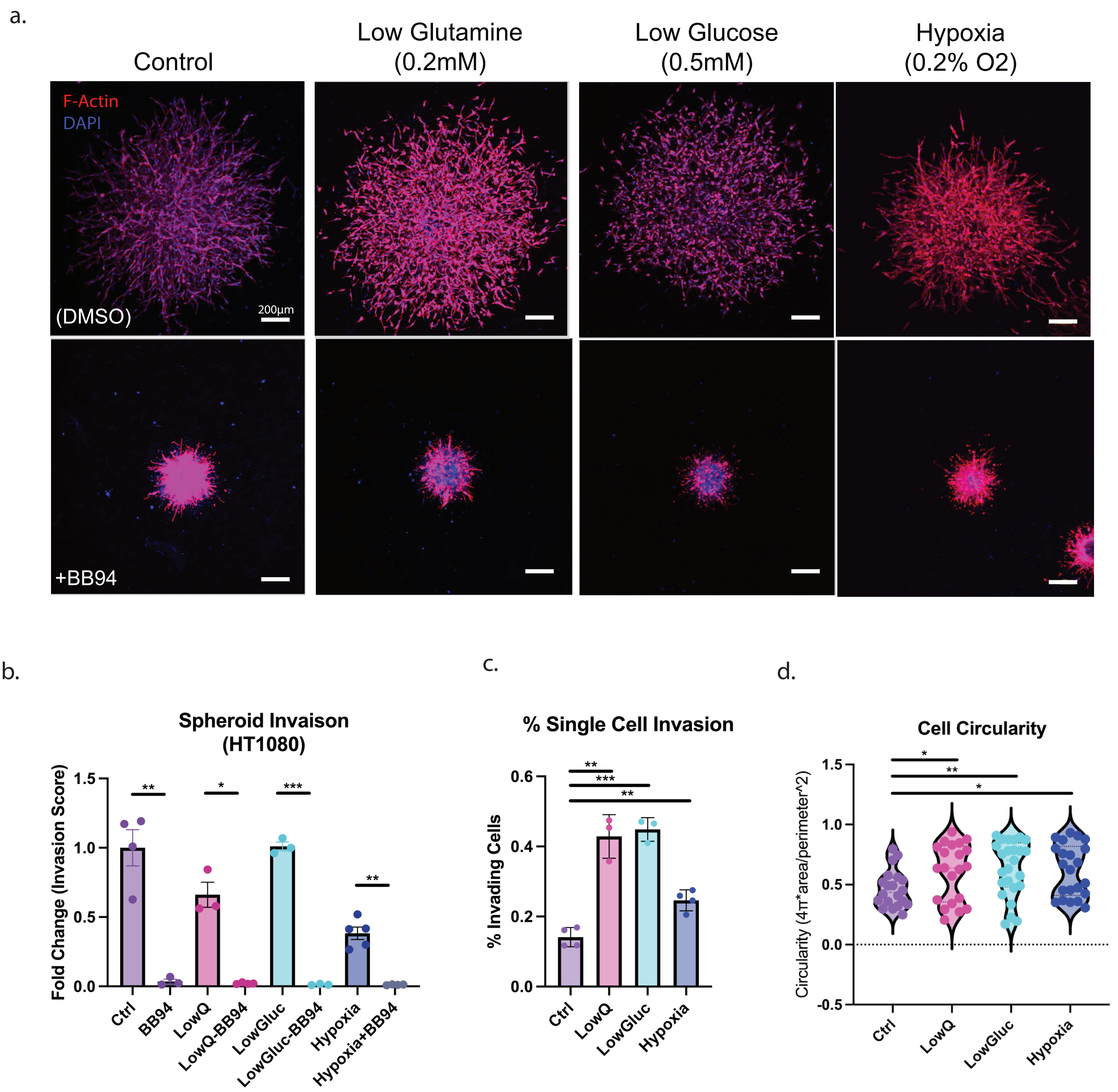


**Supplementary Figure 4: Low Nutrient-Induced Amoeboid Behavior.** a) Representative fluorescent images of HT1080 spheroids treated under the indicated conditions. b-d) Quantification of invasion, percent single-cell invasion and cell circularity for the indicated conditions. P-value < 0.05 (*), <0.01 (**), <0.001 (***), < 0.0001 (****).

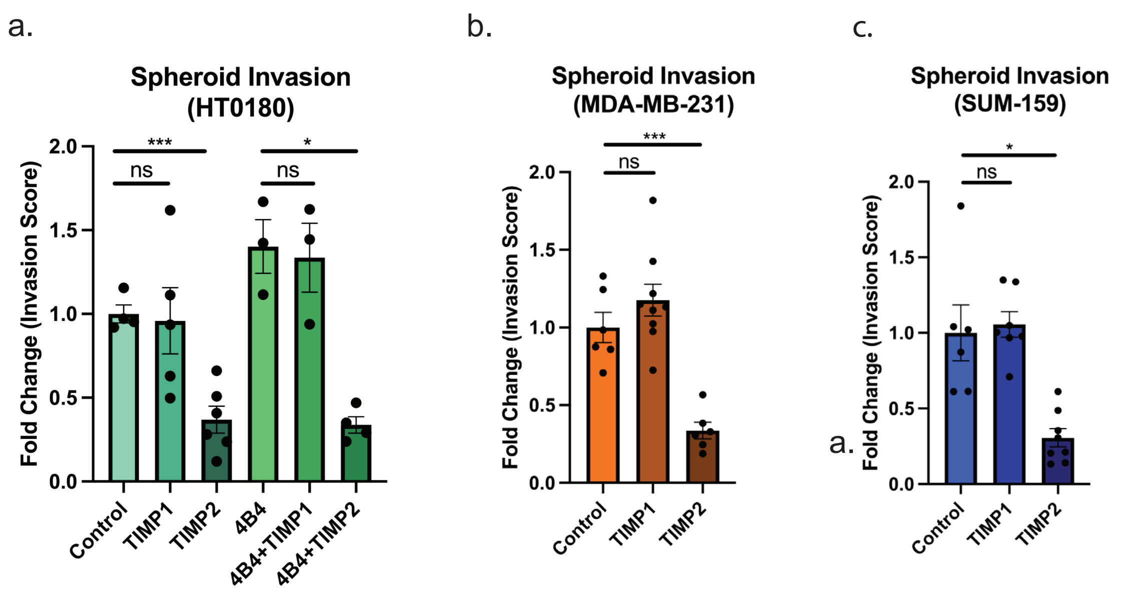


**Supplementary Figure 5: TIMP2 but not TIMP1 Disrupts Mesenchymal and Amoeboid Invasion.** a-c) Quantification of spheroid invasion in the presence/absence of either TIMP1 (12.5μM) or TIMP2 (5μM). P-value < 0.05 (*), <0.01 (**), <0.001 (***), < 0.0001 (****).

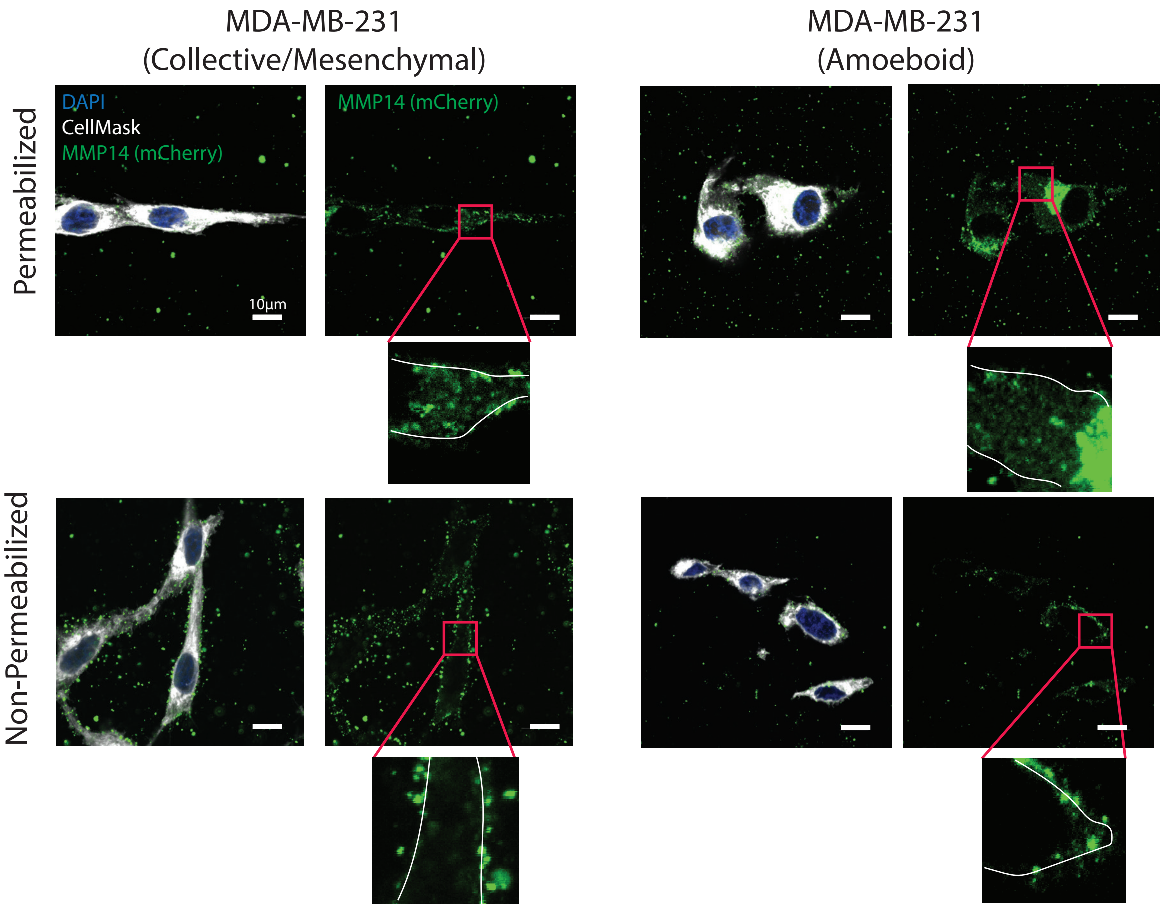


**Supplementary Figure 6: Imaging of MMP14-mCherry-KI in MDA-MB-231 Cells.** Representative images of mesenchymal and amoeboid MDA-MB-231 cells with MMP14-mCherry-KI stained with an mCherry antibody with or without permeabilization. Results representative of two separate experiments performed.

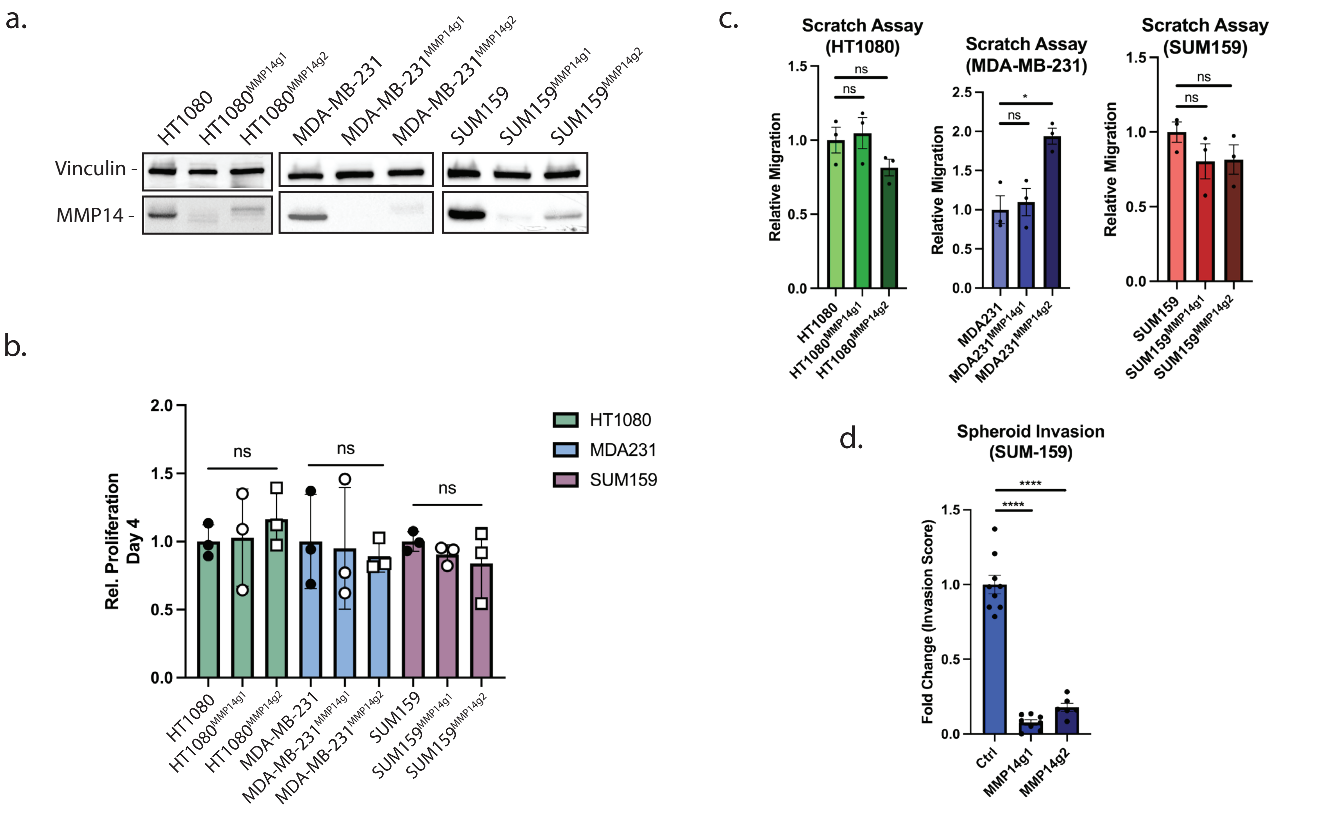


**Supplementary Figure 7: Characterization of MMP14-deleted Cells.** a) Western blot of cell lines targeted with one of two MMP14 guide RNAs. b) Quantification of proliferation of control and MMP14-targeted cells. c) Quantification of scratch assays performed using control and MMP14-targeted cells. d) Quantification of 3D spheroid invasion assay using control and MMP14-targeted SUM159 cells. P-value < 0.05 (*), <0.01 (**), <0.001 (***), < 0.0001 (****).

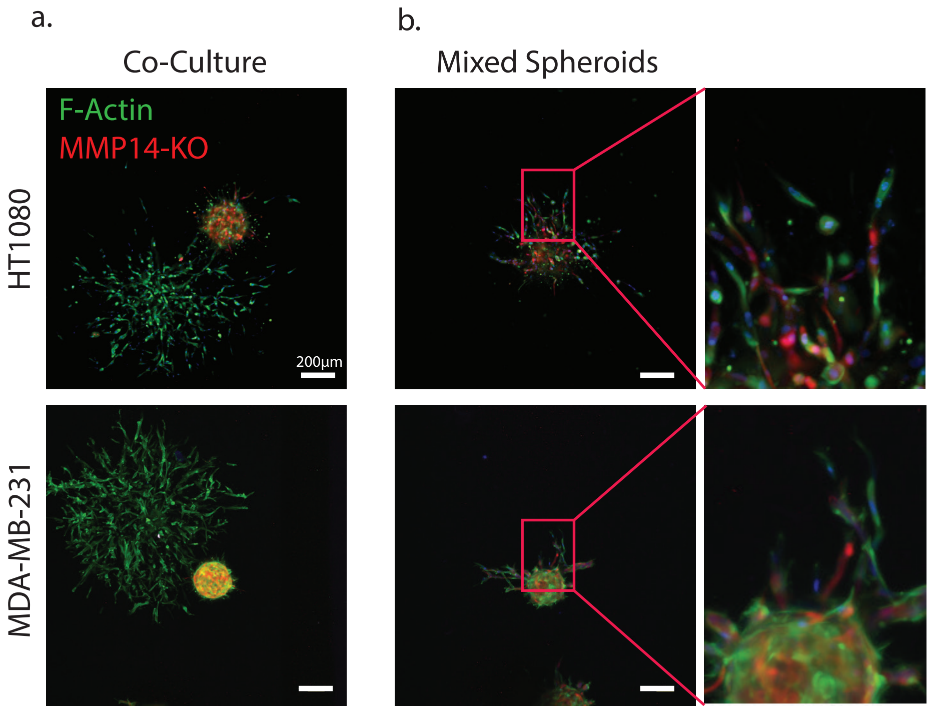


**Supplementary Figure 8: Invasive Activity of Control and MMP14-KO Mixed Spheroids.** a) Representative fluorescent images of control and MMP14-KO tumors co-cultured in close proximity. b) Representative fluorescent images of control and MMP14-KO cells mixed together within the same spheroid. Results representative of three experiments performed.

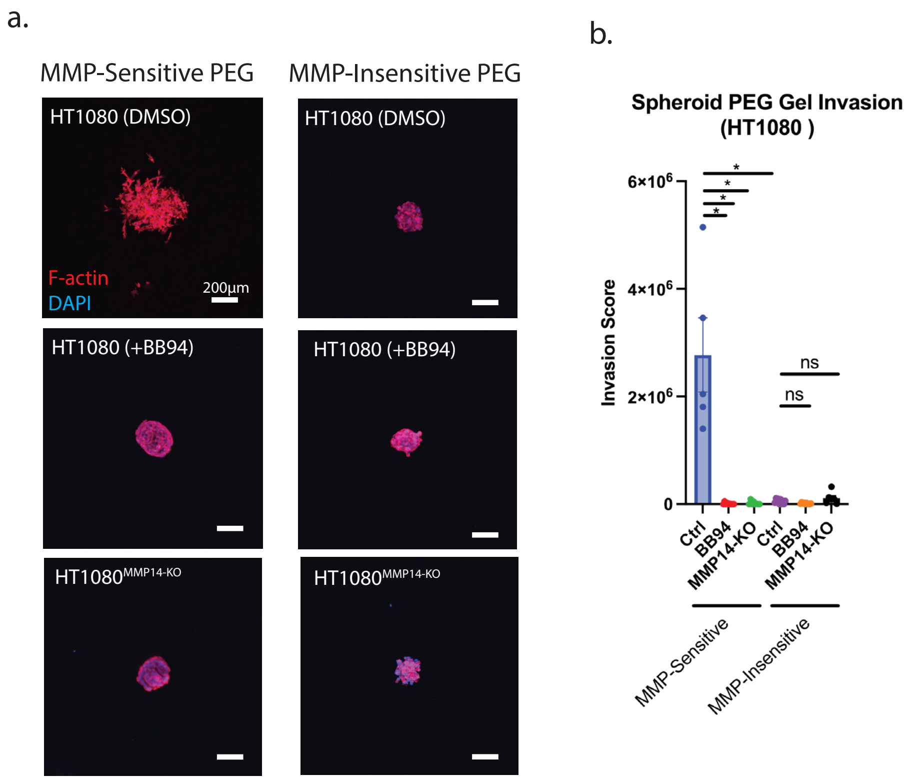


**Supplementary Figure 9: PEG Hydrogel Invasion.** a) Representative images of 4 day invasion of spheroids in either MMP-sensitive or -insensitive PEG hydrogels. b) Quantification of spheroid invasion in PEG gels. P-value < 0.05 (*), <0.01 (**), <0.001 (***), < 0.0001 (****).

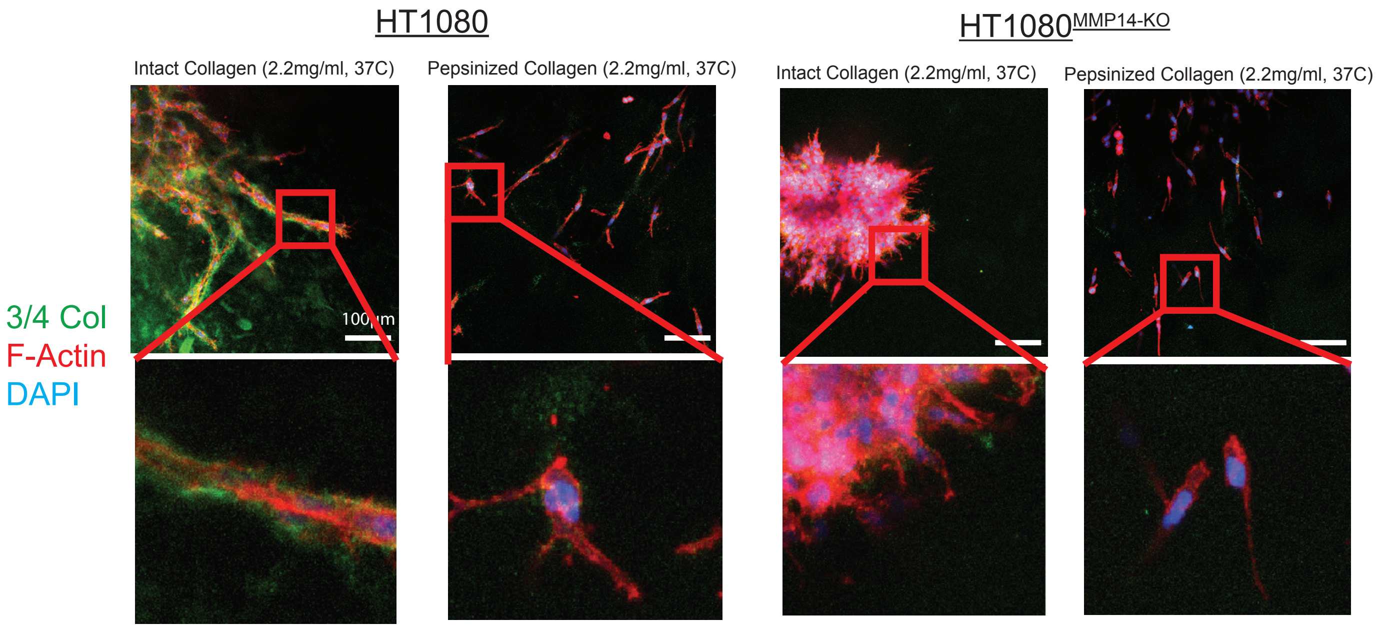


**Supplementary Figure 10: Collagen Degradation During Tumor Cell Trafficking Through Atellocollagen Hydrogels.** a) Representative images of control and MMP14-targeted HT1080 spheroids invading through 2.2mg/ml rat tail collagen versus PureCol and stained for 3/4 collagen degradation products. Results are representative of two separate experiments.

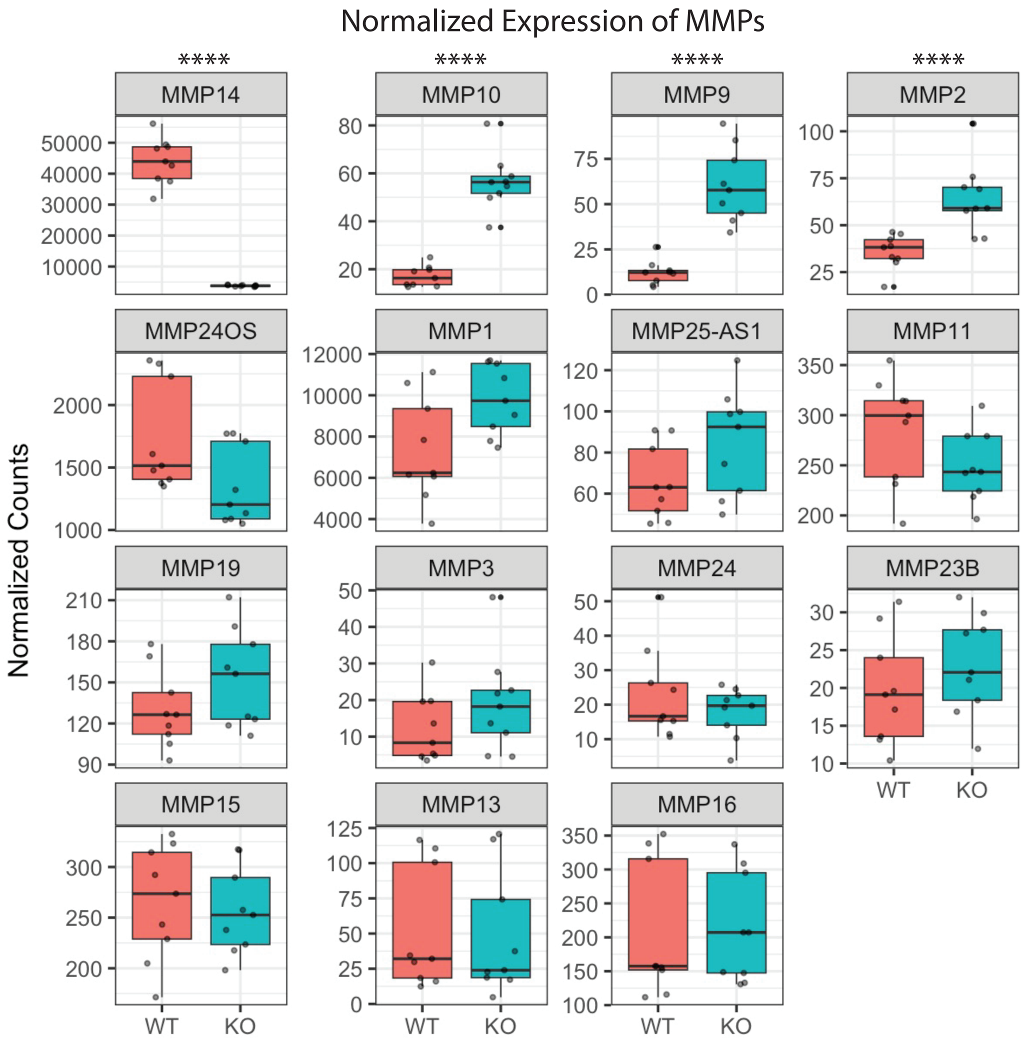


**Supplementary Figure 11: MMP Expression in MMP14-null Versus Control MDA-MB-231 Spheroids.** Boxplots showing normalized counts for MMP expression across spheroids at three timepoints (i.e., starting spheroid versus 24 and 48 -hour culture in collagen hydrogels). P-value < 0.05 (*), <0.01 (**), <0.001 (***), < 0.0001 (****).

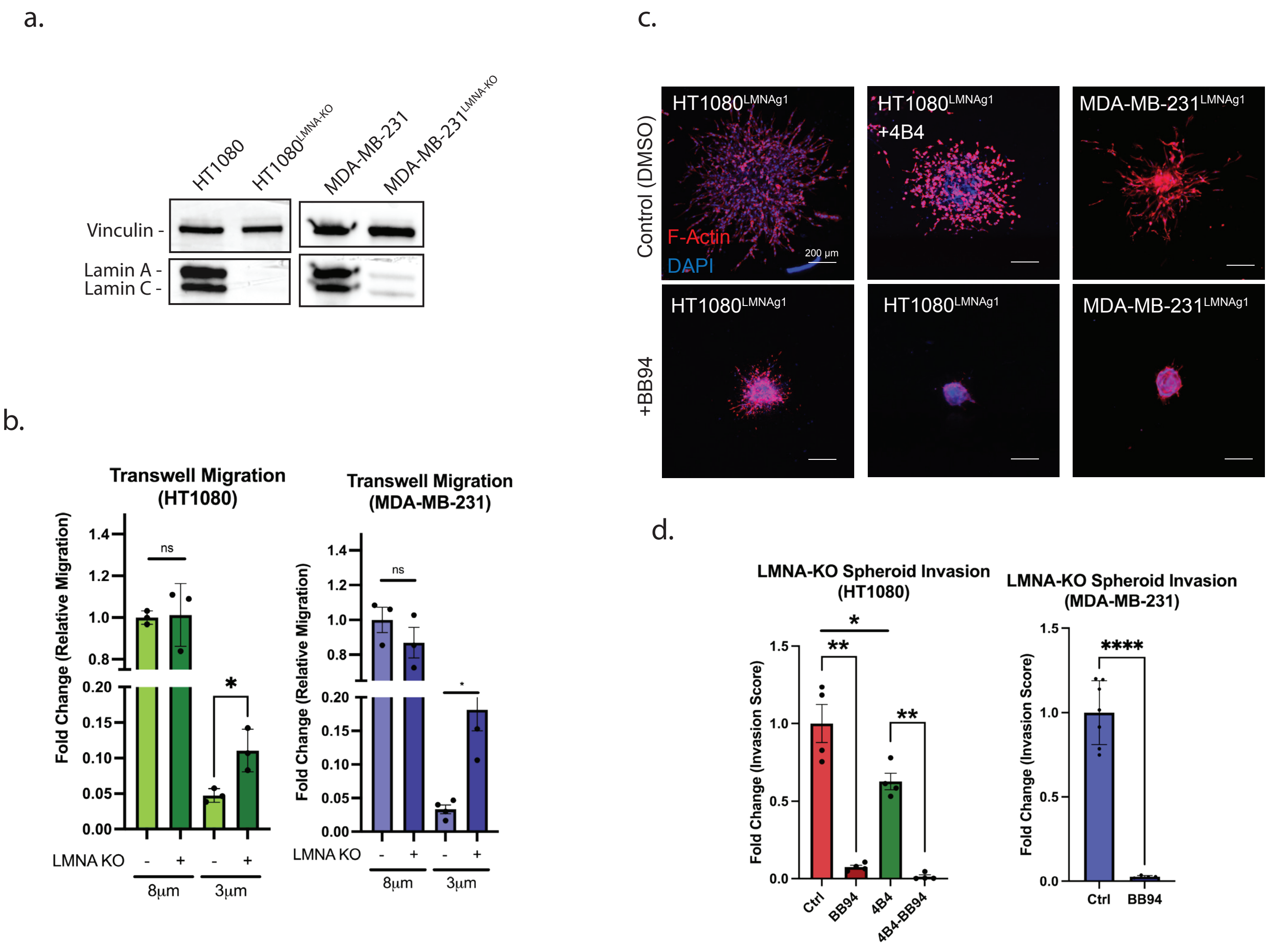


**Supplementary Figure 12: Effect of Lamin A KO on Cancer Cell Motility and Invasion.** a) Western blot of cells targeted with a LMNA guide RNA using CRISPR. b) Quantification of transwell migration assays using the indicated membrane pore sizes. c) Representative fluorescent images of spheroid invasion in 3D collagen hydrogels. d) Quantification of spheroid invasion. P-value < 0.05 (*), <0.01 (**), <0.001 (***), < 0.0001 (****).

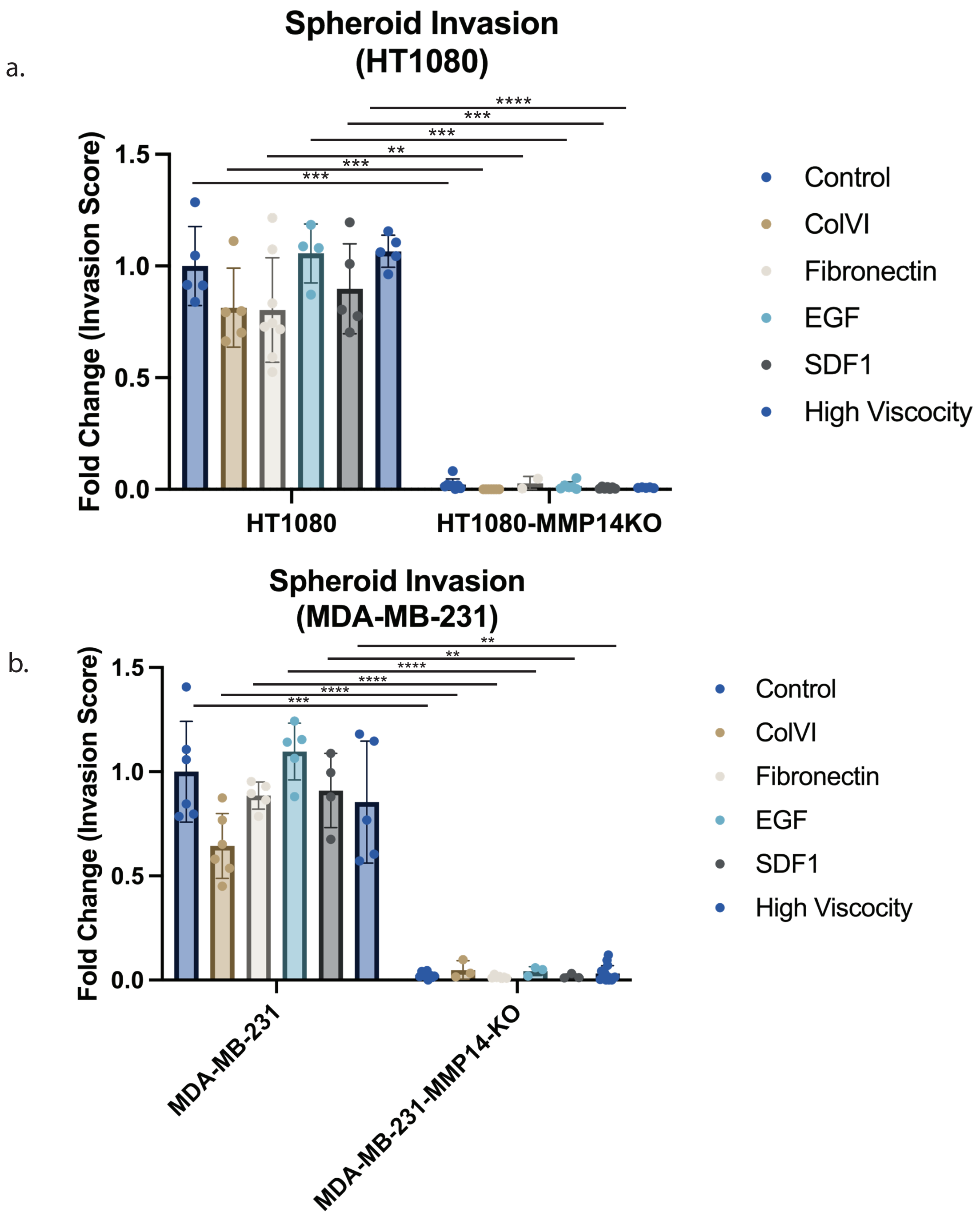


**Supplementary Figure 13: Spheroid Invasion with Hydrogel/Media Additives.** a,b) Quantification of spheroid invasion in the presence of the indicated media & hydrogel additives. P-value < 0.05 (*), <0.01 (**), <0.001 (***), < 0.0001 (****).

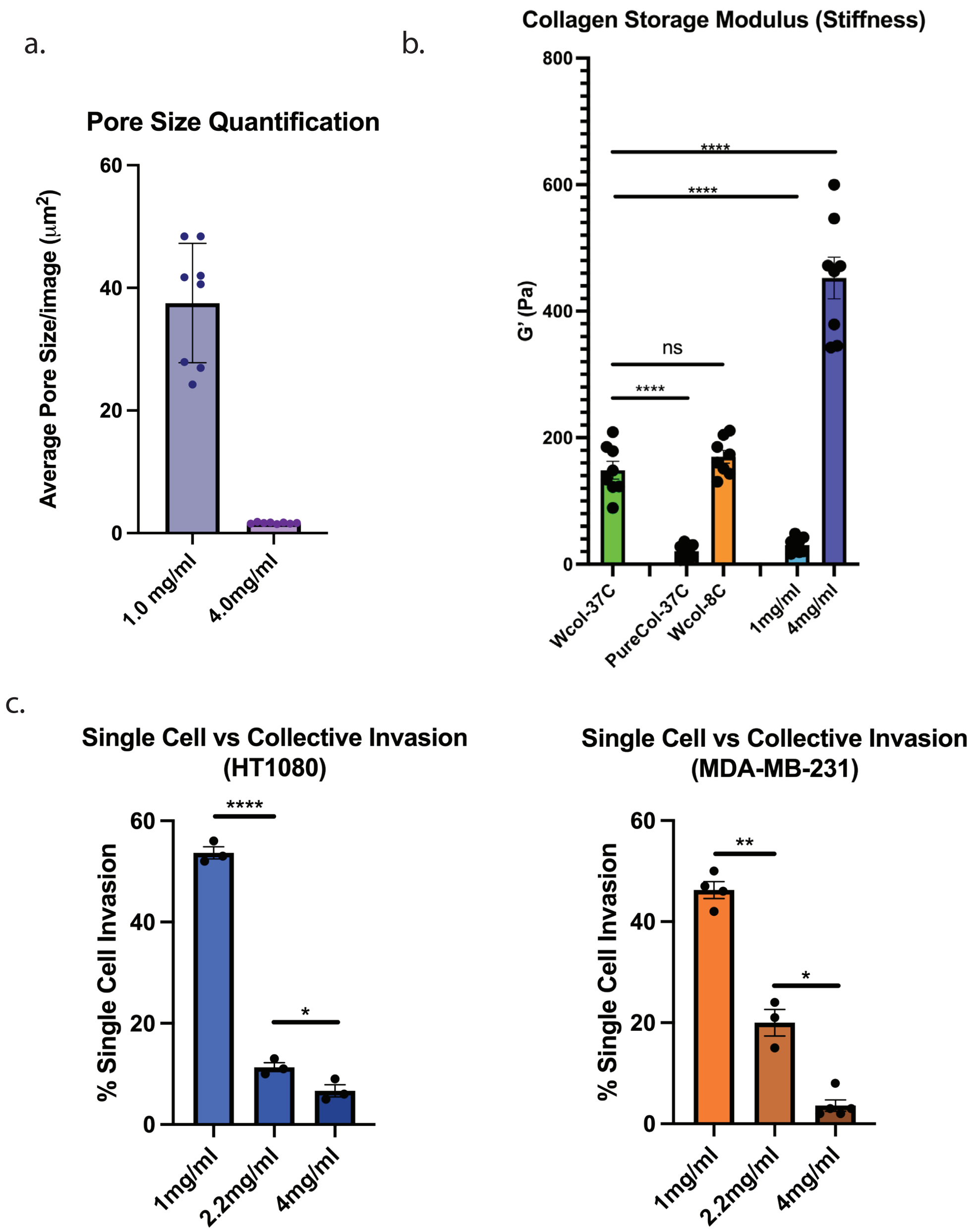


**Supplementary Figure 14: Characterzation of Hydrogel and Invasive Cell Properties with Varying Collagen Density.** a) Quantification of acid solubilized rat tail collagen pore size at the indicated concentrations. b) Quantification of storage modulus (G') for the indicated collagen hydrogel composition. c) Quantification of the percentage of single-cell invasion observed in spheroid invasion assays. P-value < 0.05 (*), <0.01 (**), <0.001 (***), < 0.0001 (****).

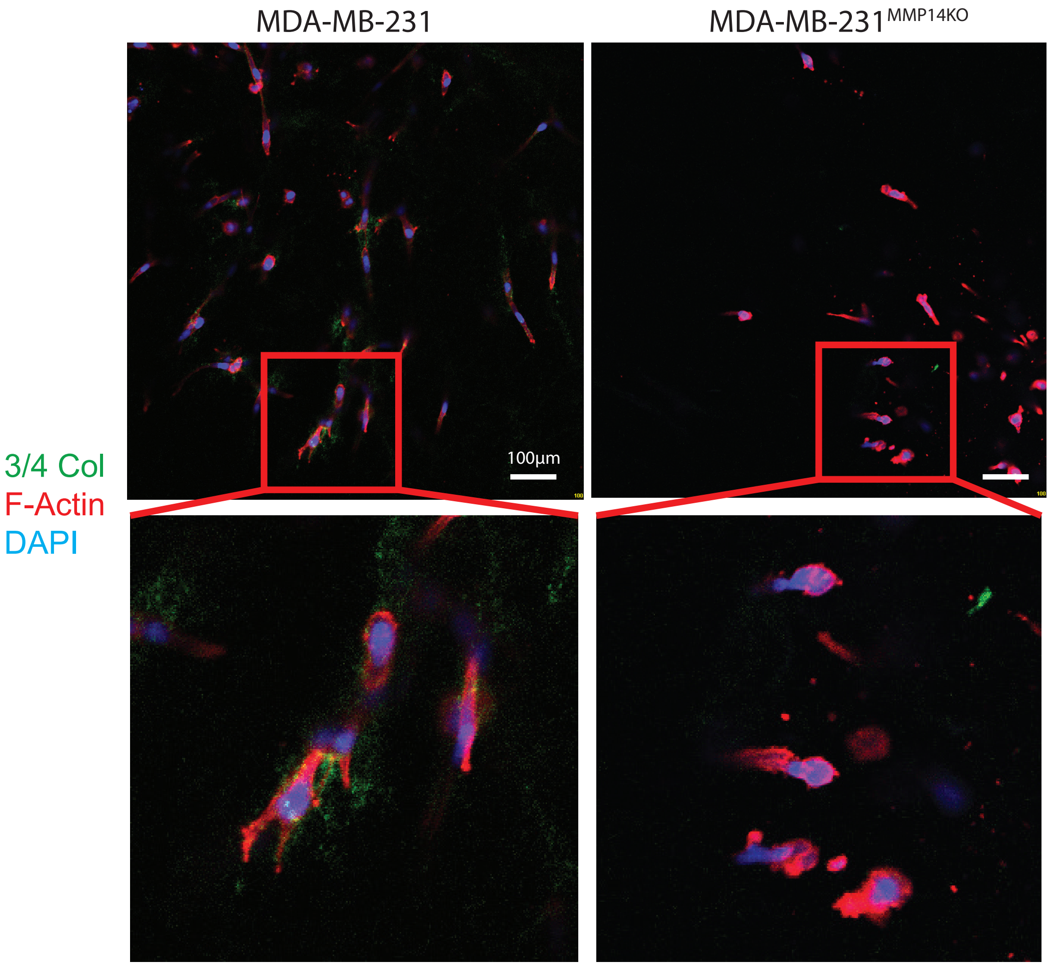


**Supplementary Figure 15: 3/4 Collagen Generation by Trafficking Cancer Cells in Low Density Collagen.** Representative immunofluorescent images of control and MMP14-KO MDA-MB-231 spheroids invading in 1mg/ml acid solubilized rat tail collagen hydrogels that were stained for 3/4 collagen degradation products. Results representative of two separate experiments.

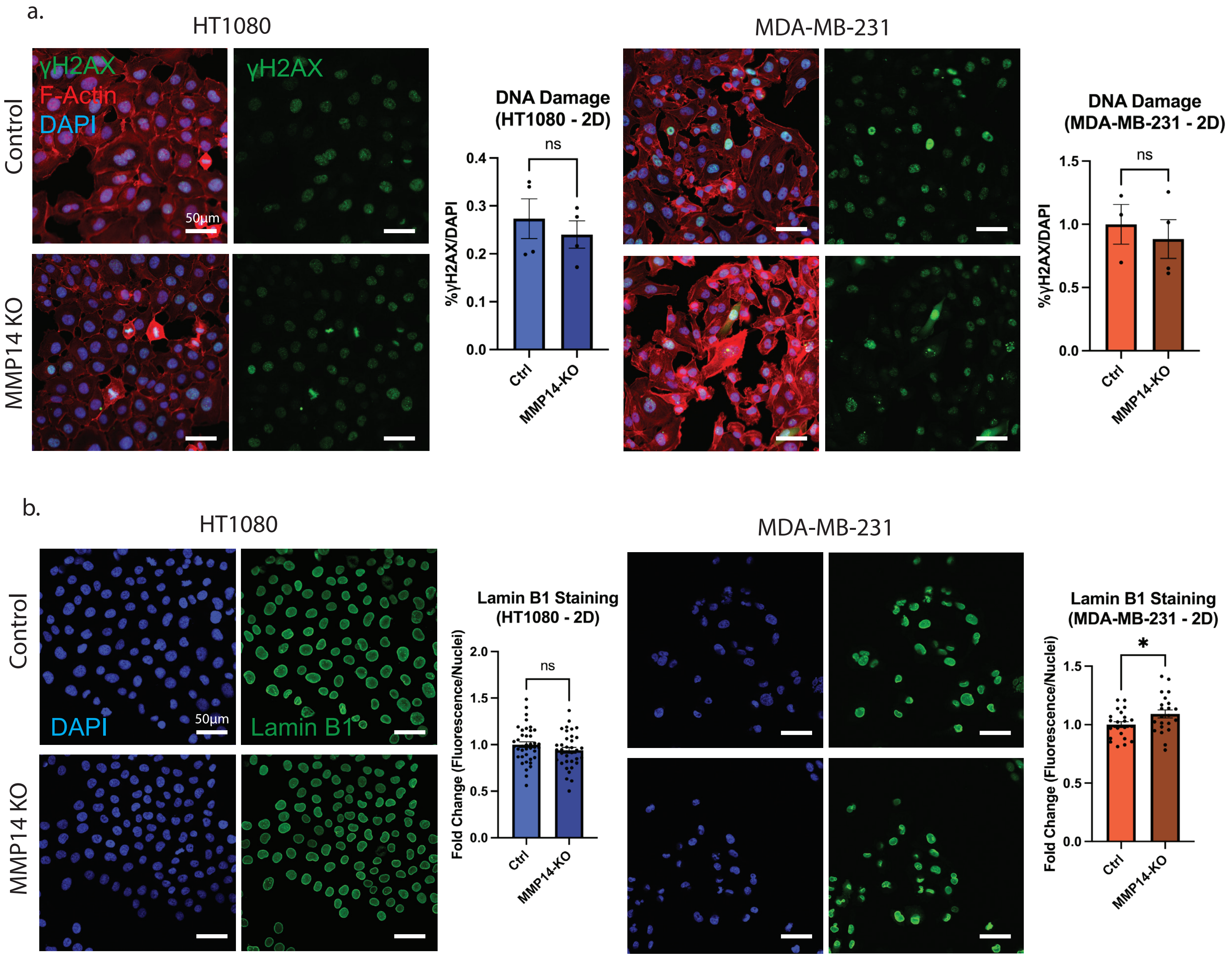


**Supplementary Figure 16: γHAX and Lamin B1 Staining in Control vs MMP14-KO Cells.**
a) Representative immunofluorescence images and quantification of control and MMP14-targeted HT1080 and MDA-MB-231 cells plated in 2D and stained for γH2AX. b) Representative immunofluorescence images and quantification of control and MMP14-targeted HT1080 and MD-MB-231 cells plated in 2D and stained for Lamin B1. P-value < 0.05 (*), <0.01 (**), <0.001 (***), < 0.0001 (****).

a.

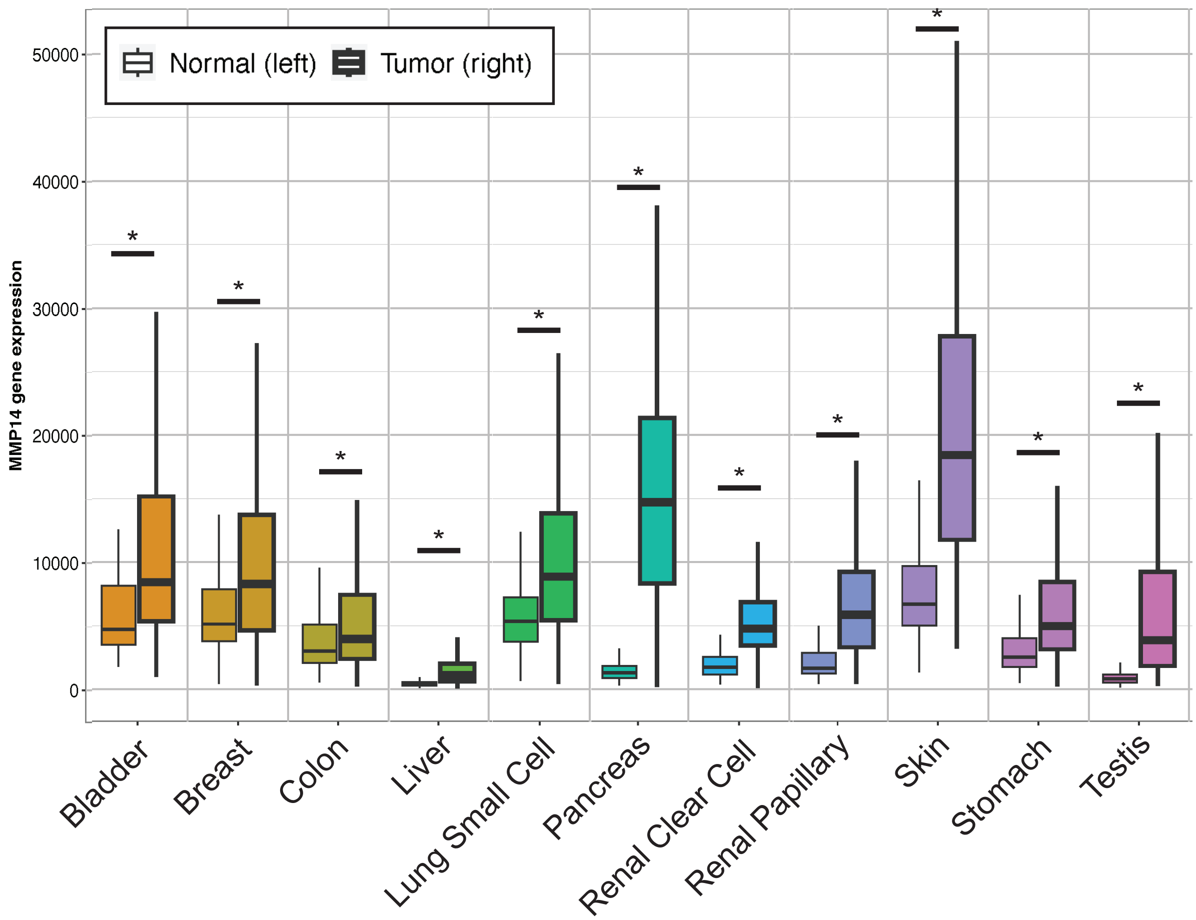


b.

| Tissue | Bladder | Breast | Colon | Liver | Lung_SC | Pancreas | Renal_CC | Renal_PA | Skin | Stomach | Testis |
|---|---|---|---|---|---|---|---|---|---|---|---|
| **Normal** | 30 | 403 | 274 | 225 | 476 | 252 | 117 | 77 | 474 | 294 | 259 |
| **Tumor** | 411 | 1097 | 469 | 371 | 501 | 177 | 535 | 289 | 103 | 375 | 156 |

**Supplementary Figure 17: Normal Versus Tumor MMP14 Expression By Tissue.** a) Box plot of MMP14 expression from RNA sequencing data including 2881 normal and 4484 tumor samples. Plot generated using TNMplot.com. b) Table displaying the number of normal and tumor samples included for each tissue type. *Mann-Whitney p<0.05 and expression >10 in tumor or normal.

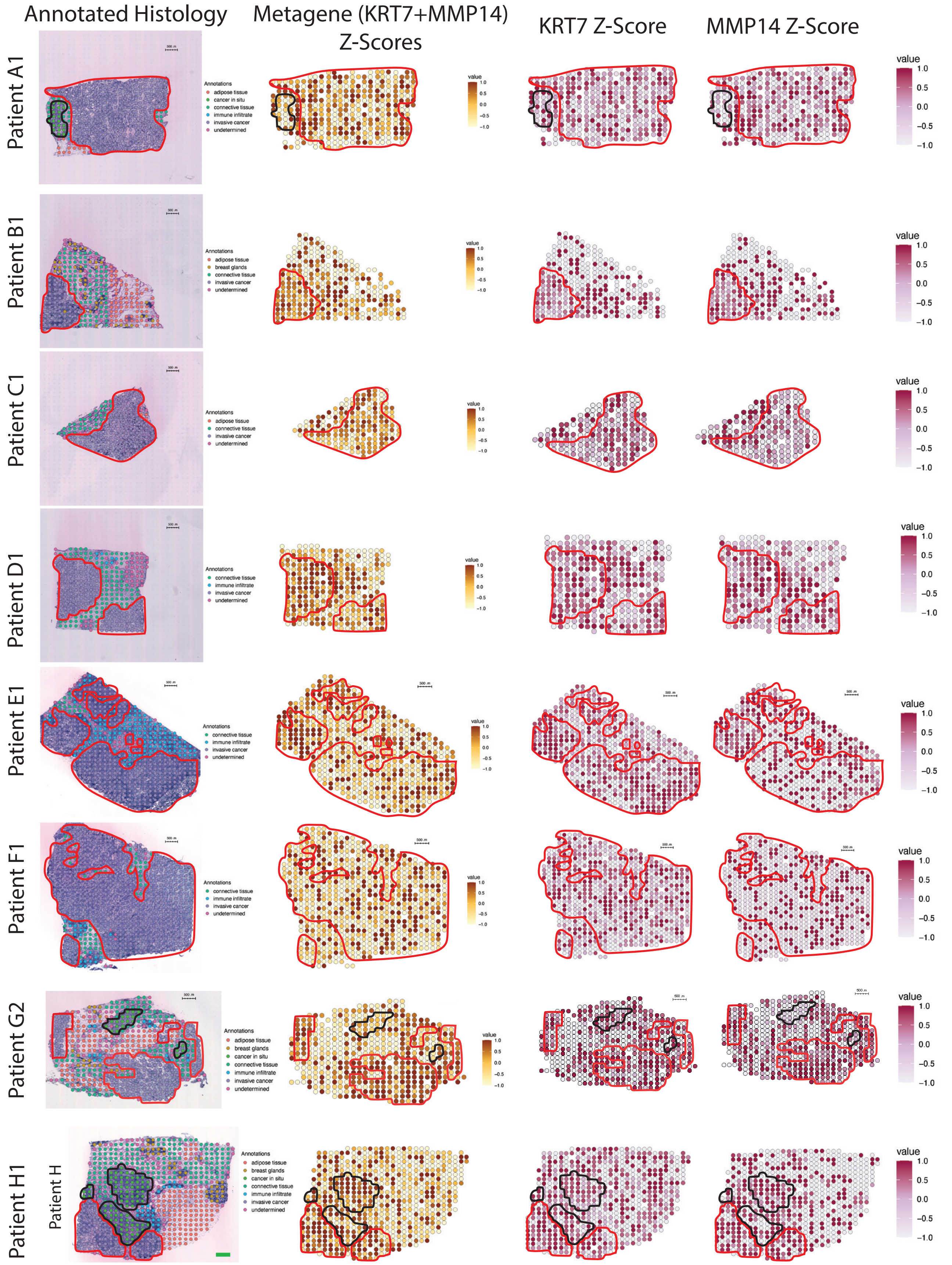


**Supplementary Figure 18: Spatial Transcriptomic Analysis of Human Breast Cancer Samples.** H&E-stained human breast cancer tissue sections with histological annotations (1st column). Z-Score spatial plots of a metagene combining normalized expression values of KRT7 and MMP14 (2nd column) as well as individual gene Z-score plots (3rd and 4th column). Red outlines indicate invasive cancer regions, black outlines indicate *in situ* cancer regions.

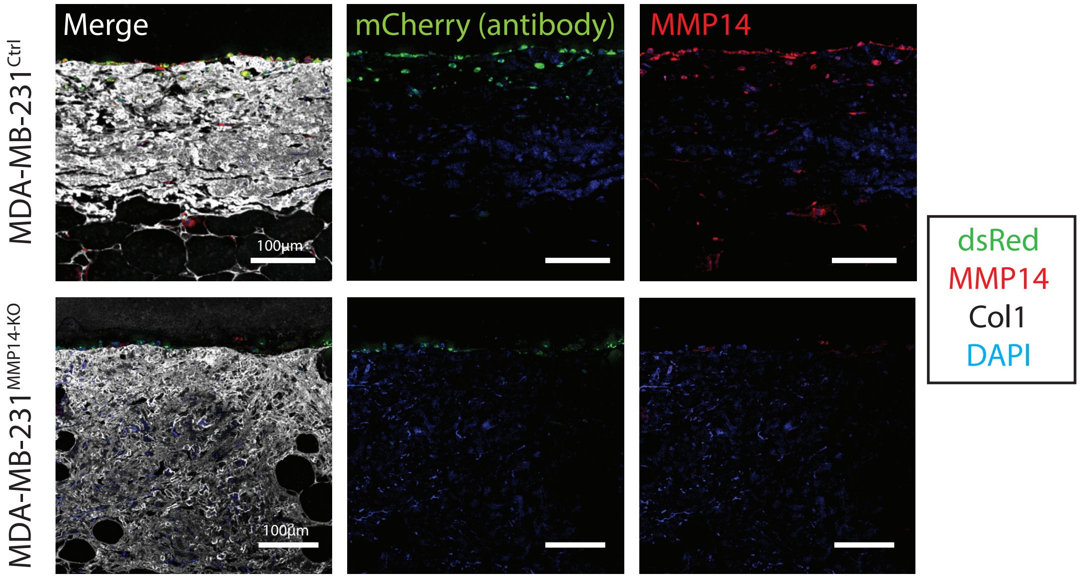


**Supplementary Figure 19: Human MMP14 Immunostaining.** Representative immunofluorescence images of control and MMP14-null MDA-MB-231 cells following 6 days of culture after plating cells atop human breast tissue in 2.5D. Staining was performed on paraffin sections and stained for the indicated antibodies. Results are representative of three separate experiments.

**Movie S1.** Phase-contrast visualization of invading control HT1080 spheroid from 24 to 48 hours post-embedding in acid solubilized rat tail collagen (2.2mg/ml, 37°C, pH 7.4).

**Movie S2.** Phase-contrast visualization of invading 4B4-treated HT1080 spheroid from 24 to 48 hours post-embedding in acid solubilized rat tail collagen (2.2mg/ml, 37°C, pH 7.4).

**Movie S3.** Spinning-disk confocal imaging of control HT1080$^{GFP-NLS}$ spheroid invasion with Trackmate tracing from 1 hour post embedding to 48 hours post-embedding in acid solubilized rat tail collagen (2.2mg/ml, 37°C, pH 7.4).

**Movie S4.** Spinning-disk confocal imaging of 4B4-treated HT1080$^{GFP-NLS}$ spheroid invasion with TrackMate tracing from 1 hour post embedding to 48 hours post-embedding in acid solubilized rat tail collagen (2.2mg/ml, 37°C, pH 7.4).

**Movie S5.** Phase-contrast visualization of invading control HT1080 spheroid from 1 hour to 72 hours post-embedding in acid solubilized rat tail collagen (2.2mg/ml, 37°C, pH 7.4).

**Movie S6.** Phase-contrast visualization of invading BB94-treated HT1080 spheroid from 1 hour to 72 hours post-embedding in acid solubilized rat tail collagen (2.2mg/ml, 37°C, pH 7.4).

**Movie S7.** Phase-contrast visualization of invading control MDA-MB-231 spheroid from 1 hour to 72 hours post-embedding in acid solubilized rat tail collagen (2.2mg/ml, 37°C, pH 7.4).

**Movie S8.** Phase-contrast visualization of invading BB94-treated MDA-MB-231 spheroid from 1 hour to 72 hours post-embedding in acid solubilized rat tail collagen (2.2mg/ml, 37°C, pH 7.4).

**Movie S9.** Phase-contrast visualization of invading MMP14-KO HT1080 spheroid from 1 hour to 72 hours post-embedding in acid solubilized rat tail collagen (2.2mg/ml, 37°C, pH 7.4).

**Movie S10.** Phase-contrast visualization of invading MMP14-KO MDA-MB-231 spheroid from 1 hour to 72 hours post-embedding in acid solubilized rat tail collagen (2.2mg/ml, 37°C, pH 7.4).

**Movie S11.** Spinning-disk confocal imaging of control MDA-MB-231$^{GFP-NLS}$ spheroid invasion from 24 to 48 hours post-embedding in low-density acid solubilized rat tail collagen (1.0mg/ml, 37°C, pH 7.4).

**Movie S12.** Spinning-disk confocal imaging of BB94-treated MDA-MB-231$^{GFP-NLS}$ spheroid invasion from 24 to 48 hours post-embedding in low-density acid solubilized rat tail collagen (1.0mg/ml, 37°C, pH 7.4).
